# Scalable generation of human stem cell-derived ureteric bud and collecting duct organoids in stirred bioreactors

**DOI:** 10.64898/2026.09.27.754790

**Authors:** Kayla J. Wolf, Jonathan E. Rubins, Ronald van Gaal, Kyle Stevenson, Aline Klaus, Pooja Nair, Jennifer A. Lewis

**Author notes:** equal contributions (Kayla J. Wolf, Jonathan E. Rubins).

## Abstract

Emerging efforts to engineer multicellular kidney tissues for drug discovery, disease modeling, and therapeutic repair require scalable generation of kidney organoids derived from human induced pluripotent stem cells (hiPSCs). Here, we report a method for generating kidney ureteric bud and collecting duct epithelial organoids in stirred tank bioreactors. We demonstrate that extracellular matrix microcarriers facilitate organoid expansion and are compatible with suspension bioreactor methods. The combination of cell-coated matrix microcarriers with stirred bioreactors promotes maturation marker expression compared to static controls comprising matrix-embedded organoids. Our method offers promise for generating epithelial organoids that serve as multicellular models and organ building blocks for scalable biofabrication.

## Introduction

Kidney organoids derived from human induced pluripotent stem cells (hiPSCs) are transforming preclinical research and offer promise as building blocks for engineering kidney tissues urgently needed for renal replacement therapy.^1–3^ For example, the FDA Modernization Act 2.0 allows preclinical testing using human cell- and organoid-based assays in place of animal models that are less physiologically relevant.^4^ Concomitantly, there is growing interest in biofabricating human kidney tissues and whole organs for therapeutic use.^5^ During embryonic kidney development, the ureteric bud (UB) grows and repeatedly branches into the surrounding metanephric mesenchyme (MM), eventually forming the collecting duct (CD) network.^6^ The ability to emulate nephrogenesis *in vitro* by spatially patterning MM and UB in three dimensions is the “holy grail” of kidney engineering. Methods for functionally connecting organoids are rapidly advancing, including biological self-assembly of kidney tubule segments^7–9^ as well as bioprinting kidney tissues composed of cell-laden inks^10^ and perfusable vessels.^11,12^ However, printing an entire organ requires over 100 billion cells.^13,14^ Hence, new methods for scalable generation of kidney organoid building blocks are needed for both high-throughput screening and organ engineering.

Stirred tank bioreactors (STRs) represent the gold standard for scalable cell expansion, as nutrients and oxygen are efficiently distributed across large volumes. To date, the development of suspension culture methods for epithelial organoids has largely focused on intestinal and hepatic organoids. Several groups incorporated low concentration liquid basement membrane matrix (BsM, 5-10% v/v) in medium to promote organoid suspension culture.^15–17^ Stirred or rotational culture maintained or improved organoid growth and maturation compared to conventional static culture. Alternatively, Co *et al.* rapidly generated solid BsM droplets and filaments which encapsulated multiple intestinal organoids.^18^ Suspension droplet culture improved organoid uniformity compared to static culture. Although these studies show that incorporating matrix with suspension culture can support or enhance epithelial organoids, the influence of STR conditions on cells during differentiation remains undefined.

Two types of kidney organoids can be derived from hiPSCs: (1) MM organoids that give rise to nephron structures^19,20^ and (2) UB organoids that can be further differentiated into CD organoids.^21–24^ Recently, modified protocols for producing nephron-rich organoids in STRs have been developed to improve scalability.^25–27^ Despite these recent advances, analogous methods for the scalable differentiation of UB/CD organoids have yet to be reported. We posit that two unique features of UB/CD organoid differentiation (relative to MM organoids) hinder the modification of static UB/CD differentiation protocols to STRs. First, UB/CD organoids are generated by embedding ureteric progenitor cell (UPC) aggregates within a 3D BsM, *i.e.*, Geltrex or Matrigel, to support branching morphogenesis and normal epithelial polarization.^21,28,29^ Second, the effect of stirred culture conditions on UB/CD organoids is unknown, although likely significant. For example, stirring induces fluidic shear stress, which is known to influence stem cell differentiation.^30^ As previously shown, renal epithelium is particularly sensitive to low fluidic shear stress (∼0.01 to 1 dyn/cm^2^)^31–33^ relative to other epithelial and endothelial cells.^34–36^ Hence, the successful modification of static UB/CD differentiation protocols to STRs requires understanding both matrix and stirring effects on their differentiation.

Here, we report a method for differentiating UB/CD organoids in STRs uniquely enabled by using matrix microcarriers composed of fragmented BsM that support cell growth and modulate differentiation. Fragmented microcarriers are rapidly produced from multiple ECMs. We show that the incorporation of fragmented BsM microcarriers with hiPSC-derived UPCs in STRs leads to upregulation of key maturation markers for both UB and CD organoids compared to static controls. The scalable generation of UB and CD organoids provides a foundational step toward engineering multicellular kidney tissues for drug discovery, disease modeling, and, ultimately, therapeutic repair.

## Results

### UB/CD organoids embedded in matrix via static differentiation conditions

We generated UB organoids following the protocol established by Zeng & Huang *et al.* (**Table S1**), which results in more highly expandable UB tip-like cells^21^ compared to other reported protocols.^23,28^ We began by differentiating hiPSCs to UPCs and the further differentiated these UPCs to UB organoids using two static culture formats: standard 96-well plates and 6-well plates fitted with ∼300 microwells/well inserts. For the later format, we first aggregated UPCs in commercially available 800 µm diameter microwell plates and then embedded the resulting multicellular aggregates in BsM in static cell culture inserts (24 mm diameter), with ∼300 organoids per insert fitted within a single well of a 6-well plate (**Fig. 1A**). All UB organoids expanded in size at similar rates compared to those cultured in 96-well plates (one organoid/well), although the normalized size of UB organoids cultured in inserts is slightly higher on day 35 (**Fig. 1B-C**). Morphologically, UB organoids generated under both static culture formats at each timepoint, day 21 and day 35, exhibited large bud-like structures with epithelialized, open lumens (**Fig. 1C-D**). We confirmed bud and luminal morphology for UB organoids embedded in BsM (50% vol/vol Matrigel or Geltrex) (**Fig. S1**). The UB epithelial cells expressed key UB protein markers, including SOX9, CDH1, KRT8,s ETV5, RET, GATA3, and PAX2 (**Fig. 1D, Fig. S1**).

**Figure 1.**
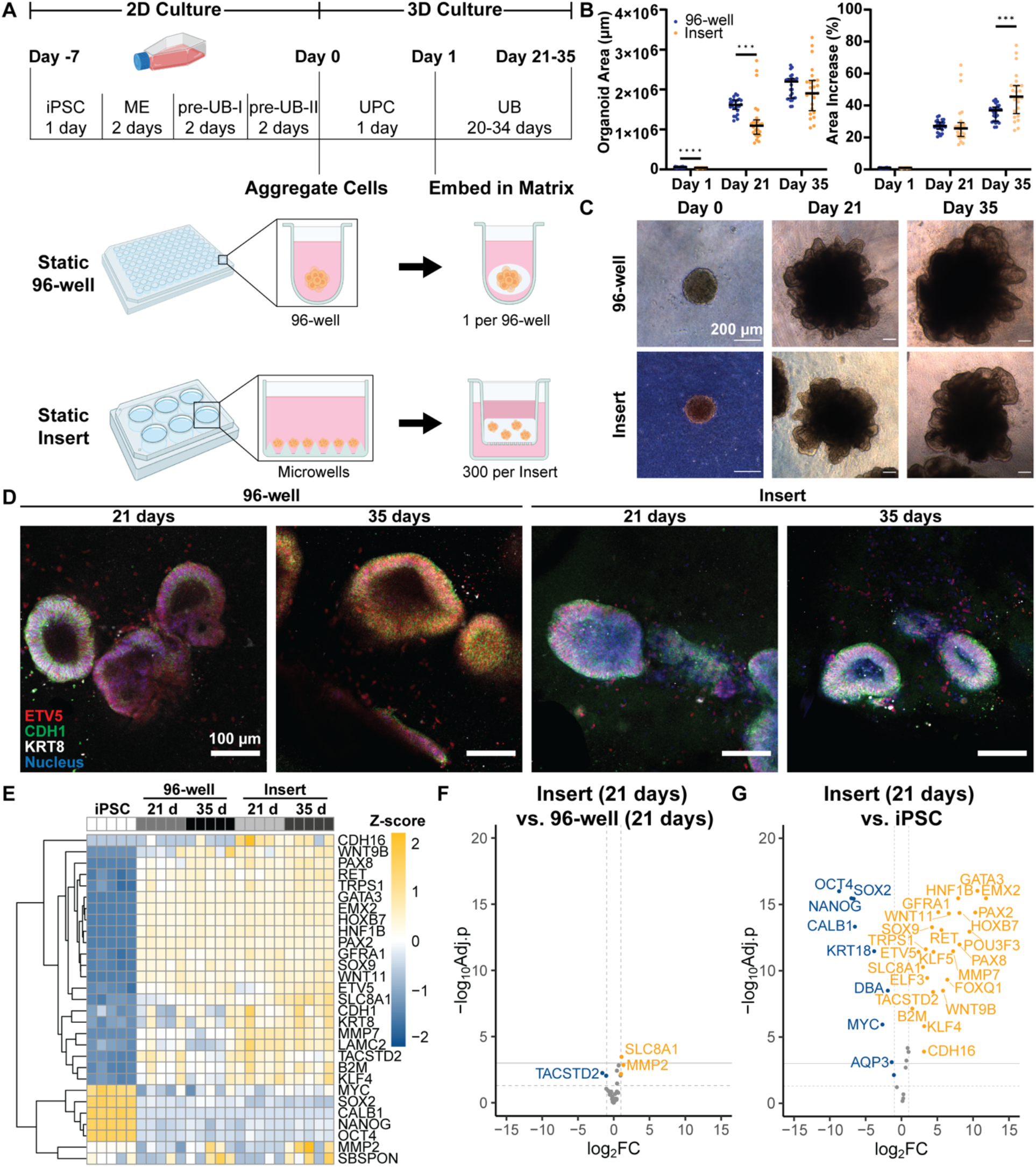
UB organoid generation. A) Overview of UB differentiation protocol adapted from Zeng & Huang, et al.^21^. hiPSCs are cultured in 2D flasks from day -7 to day 0. On day 0, cells are aggregated in microwell plates. On day 1, aggregates are embedded in BsM in static cell culture inserts. UB organoids are cultured in UBCM to day 21 or day 35. B) Organoid area over time in 96-well plates compared to inserts. Left = raw measurements, right = normalized to day 1. N = 45 organoids (dots) from 3 independent differentiations. Bars represent median and interquartile range. ***p<0.001, ****p<0.0001 using unpaired t-tests with two-stage step-up false discovery rate (FDR) correction. C) Brightfield images demonstrating organoid area and morphology over time in 96-wells compared to inserts. Scale = 200 μm. D) Immunofluorescence images of tubules in 96-well and insert organoids at 21 days and 35 days. Red = ETV5, green = CDH1, white = KRT8, blue = nucleus. Images are single slices. Scale = 100 μm. E) Heatmap of selected genes in hiPSCs compared to organoids cultured in 96-well or insert plates for 21 and 35 days. N = 5 independent differentiations. Colors correspond to up- (orange) and down-(blue) regulated genes scaled to the standardized *Z* score. F) Expression of UB, CD, and stemness genes in 21-day old organoids cultured in inserts compared to iPSCs. G) Gene expression in 21-day old organoids cultured in inserts relative to 21-day old organoids cultured in 96-well plates. For (F) and (G), all tested genes above threshold are shown with horizontal dashed line marking adj. p<0.05, horizontal solid line marking adj. p<0.01, and vertical dashed line marking 2-fold change. Data points represent individual genes from N = 5 independent experiments, where *p* values are adjusted by the Benjamini-Hochberg correction.

We find that the UB organoid transcriptome is similar regardless of static culture format or time (**Fig. 1E-F Fig. S2**). Our panel included 172 genes that comprise known markers of UB, CD, MM, stemness, and off-target cell types (*e.g.*, neuronal markers^37^), as well as genes for ECM components (**Table S2**). As expected, the UB organoids exhibited increased UB and CD marker expression generally (*e.g*., *PAX2, GATA3, WNT9B, WNT11, RET*) and decreased stemness marker expression (*e.g*., *OCT4, SOX2, NANOG, MYC*) relative to hiPSC controls indicating differentiation. (**Fig. 1G, Fig. S2**). We recognize that some markers may be imprecisely expressed or have multiple roles in development, thus reflecting both an iPSC-state as well as a terminally differentiated state. We see, for example, that *CALB1* (a known CD marker) is higher in hiPSCs than in UB (**Fig. 1G, Fig. S2**). Several genes (*e.g., TACSTD2* and *MMP2*) are significantly differentially expressed in the insert-cultured UB organoids relative to 96-well controls (**Fig. 1F, Fig. S2**); however, the observed fold changes in those genes are minimal (<1.5).

Next, we used established protocols to further differentiate UB organoids embedded in BsM to CD organoids^21,24,28^ in cell culture inserts (∼300 organoids per insert fitted within a single well of a 6-well plate) under static conditions. We find that key markers of UB stalks and mature CD (e.g., *CALB1, WNT9B, WNT7B*) are upregulated, while UB tip markers (*RET, WNT11, ETV5*) are downregulated (**Fig. S3**). Immunofluorescent staining confirms decreased RET expression at the protein level in CD organoids compared to UB organoids (**Fig. S4**). By contrast, there are no clear changes in KRT8 expression at the gene or protein level **(Figs. S3-S4**). The CD organoids are dominated by AQP2^+^ principal-like cells with primary cilia (**Fig. S5A**). Consistent with others^21^, we did not see evidence of intercalated cells in these organoids (**Fig. S5B**). To further improve scalability of the protocol, we eliminated the CD117^+^ cell sorting step and observed no significant differences in morphology or transcriptome between CD117^+^-derived organoids and unsorted UB organoids (**Fig. S6**). Hence, one can scale UB and CD organoid differentiation from 96-well plates to multi-well cell culture inserts for use from 21-35 days old.

### Matrix microcarriers for UB and CD organoid differentiation

Existing UB differentiation protocols are not amenable to culture in STRs as they require embedding within a three-dimensional matrix. To enable STR culture, we explored two approaches of aggregating cells with BsM: (1) droplet encapsulation and (2) fragment-based microcarriers (**Fig. 2A**). In exploratory experiments, we emulated the STR microenvironment by culturing UB organoids in suspension using 6-well plates and T25 flasks placed on orbital shakers. In the first approach, we manually formed droplets by encapsulating cell aggregates composed of UPCs in 100% Geltrex via the hanging drop method (**Figs. S7-S8**).^38^ We note that droplet encapsulation can also be automated via droplet microfluidics.^39^ In the second approach, microcarriers composed of 100% Geltrex fragments, previously show to range from 500-8,500 µm^2^ (2D area) ^40^, are produced by extrusion fragmentation (**Fig. S7**). These fragments are then mixed with dissociated UPCs that adhere to the outer surfaces of individual fragments overnight. These two methods result in different configurations, *i.e*., UPC aggregates encapsulated within droplets that expand towards and along the outer droplet surface, while UPCs adhered to and expanded on fragment surfaces (**Fig. 2B**). We typically observed a complex internal tubular architecture akin to organoids cultured in droplets which we posit forms by the aggregation of multiple fragments (**Fig. 2C-D**). The tubular lumens inside the fragment-based organoids are contiguous with the outer surface, enabling primary cilia within the lumens to be directly exposed to culture medium (**Fig. 2D, Movie S1**). Notably, the droplet approach enables the organoid tubular network to grow inside-out while the fragment approach simultaneously facilitates outside-in and inside-out network formation (**Fig. 2B-D**).

**Figure 2.**
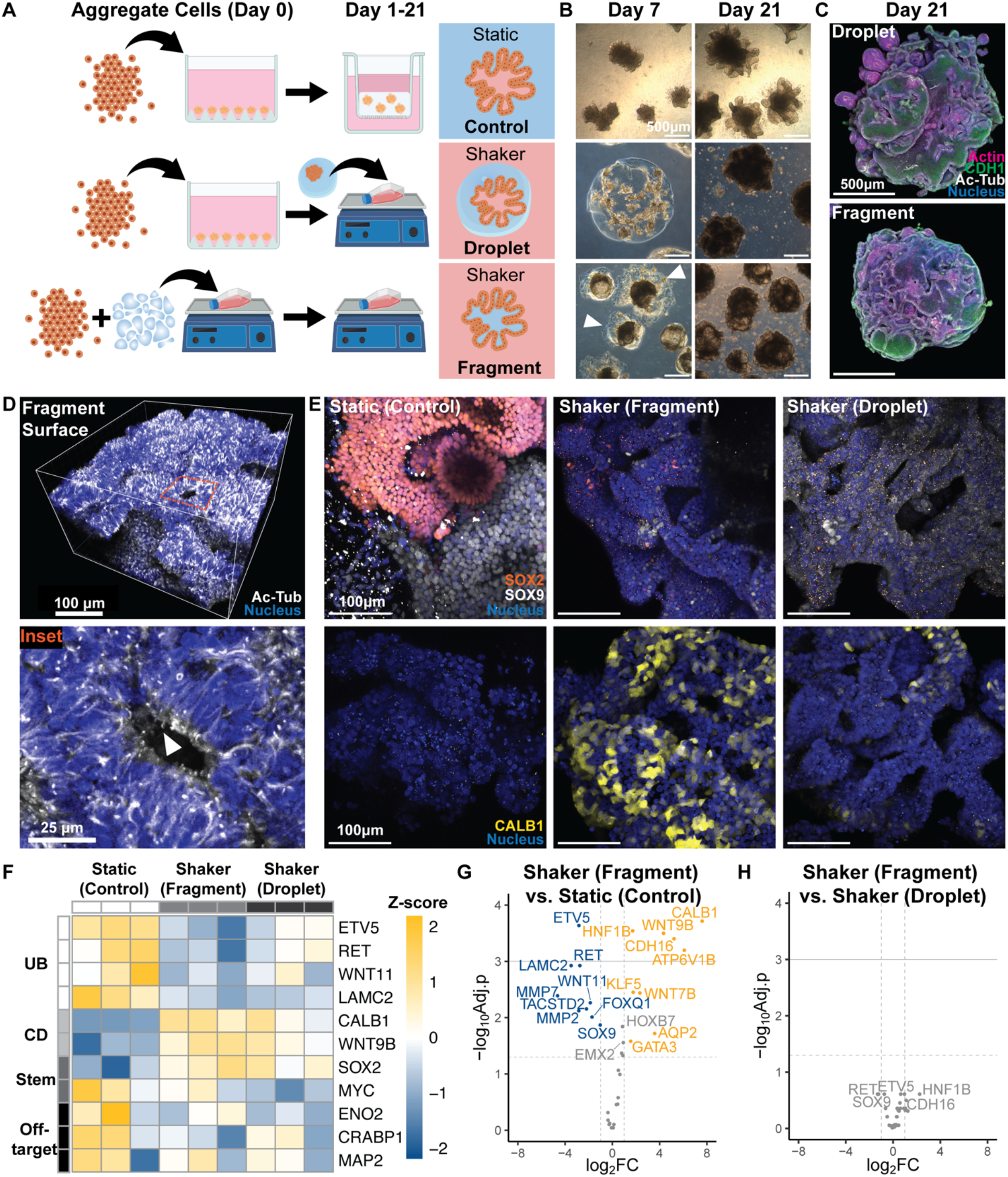
Matrix microcarriers for UB organoid culture on orbital shakers. A) Overview of culture workflow. UPCs are either combined with fragmented BsM or aggregated and embedded in BsM droplets, then cultured in a non-adherent T25 flask on an orbital shaker. B) Morphology of UB organoids (shaker and static control) after 7 and 21 days in culture. White arrows indicate fragment surface area not yet overgrown with cells. Scale = 500 μm. C) Maximum intensity projections showing internal structures of organoids encapsulated in BsM droplets compared to those cultured with BsM fragments after 21 days in culture, both cultured on orbital shakers. Magenta = actin, green = CDH1, white = Ac-Tub, blue = nucleus. Scale = 500 μm. D) Perspective view of organoid surface cultured on fragment microcarriers after 21 days in culture. Orange box indicates inset. White = Ac-Tub, blue = nucleus. Scale = 100 μm. Inset shows apically-oriented cilia in lumen that is contiguous with surface. White arrow marks a primary cilium. Scale = 25 μm. E) Maximum intensity projections of key UB markers for organoids at 21 days in culture. Orange = SOX2, white = SOX9, yellow = CALB1, blue = nucleus. Scale = 100 μm. F) Heatmap of key markers for UB, CD, stemness (stem), and off-target neuronal populations in each culture condition. N = 3 independent differentiations. Colors correspond to up-(orange) and down-(blue) regulated genes scaled to the standardized *Z* score. G) Gene expression in organoids cultured on fragment microcarriers on an orbital shaker relative to static embedded controls. H) Gene expression in shaker organoids cultured on fragment microcarriers relative to organoids embedded in droplets. For (G) and (H), only UB and CD genes tested above threshold are shown. Horizontal dashed line marks adj. p<0.05, horizontal solid line marks adj. p<0.01, vertical dashed line marks 2-fold change. Data points represent individual genes from N = 3 independent experiments, where *p* values are adjusted by the Benjamini-Hochberg correction.

As a benchmark, we compared the resulting morphology and gene expression of UB organoids produced by the droplet encapsulation and matrix fragmentation methods to those generated via static embedded culture in multi-well cell culture inserts and to control organoids cultured without matrix (no BsM) in the same manner. Immunofluorescent staining of several key markers demonstrated differential expression of SOX2, SOX9, and CALB1 at the protein level (**Fig. 2E, Fig. S9**). While large clusters of SOX2^+^ cells are present in static conditions and matrix-free conditions, they are largely absent in UB organoids produced on an orbital shaker via droplet encapsulation or matrix fragmentation (**Fig. 2E, Fig. S9**). Stemness is clearly reduced in UB organoids subjected to fluid flow in the presence of BsM. Transcriptomic analysis revealed that UB organoids generated via shaker culture with BsM exhibited downregulation of UB tip marker expression (*e.g*., *ETV5, RET, WNT11*) and upregulation of stalk and maturation markers (*e.g*., *CALB1, WNT9B*) compared to those in static culture (**Fig. 2F-G, Fig. S10**). Notably, there were no significant differences in gene expression between droplet encapsulated and fragment-based organoids (**Fig. 2H**). Morphologically, all organoids containing matrix are more homogenous and contain fewer non-lumenized spheroids (**Fig. 2E, Fig. S9**). By contrast, both BsM-free controls (static and suspension) exhibit highly heterogeneous morphologies, including round and elongated clusters of cells (**Fig. S9**).

We then investigated whether aggregated hiPSCs, termed embryoid bodies (EBs), could be used directly with matrix fragments on orbital shakers to generate UB organoids (**Figs. S11-S12**). Compared to matrix-free controls, we observed that matrix fragmentation coupled with fluid flow (shaking) enhances the morphological homogeneity of these UB organoids compared to highly heterogeneous UB organoids cultured without matrix (**Fig. S11B**). Likewise, transcriptomic analysis reveals upregulation of many UB/CD genes (primarily stalk and some tip markers) and downregulation of stemness markers (*e.g*., *KLF4, MYC, SOX2*) and off-target neuronal markers (*e.g*., *CRABP1, ENO2, MAP2*) in fragment-based organoids derived directly from hiPSCs (**Fig. S11C-E, Fig. S12**). Immunofluorescent staining confirms expression of several key UB markers (*e.g*., CDH1, KRT8, RET) as well as epithelial-like, cobblestone morphology in EB-derived organoids relative to controls, thereby resembling trends observed in UPC-derived organoids (**Fig. S11F**).

Importantly, these exploratory experiments reveal that UB organoids can be differentiated from both UPCs and iPSCs in shaker culture leading to an enhanced transition from tip-like, immature UB cells to more stalk-like, mature UB cells, and that matrix microcarriers aid this transition. While both droplet and fragment microcarriers increased UB maturation compared with static embedded controls, the fragmentation approach is more amenable to further scale-up using stirred bioreactors, since milliliters of fragments can be produced within minutes. Moreover, the complex internal architecture of UB organoids differentiated on matrix fragments may provide additional benefits such as enhanced nutrient diffusion through open lumens. Notably, SOX2^+^ clusters of cells were observed in the static embedded organoids, but not in the EB-derived shaker organoids with or without matrix (**Fig. S11G**). These results collectively suggest that deriving UB organoids from EBs is feasible, and that differentiation is enhanced by the inclusion of matrix microcarriers.

### Scalable UB and CD organoid differentiation via matrix microcarriers in stirred bioreactors

As a final demonstration, we generated UB organoids by co-introducing UPCs (day 0) or hiPSCs (EBs, day -7) with fragmented matrix microcarriers in STRs (**Fig. 3A, Fig. S13A**). After 21 days, we observed epithelial morphology under phase contrast in all conditions, although organoids derived from EBs are more heterogenous and contained dark clusters of mesenchymal cells (**Fig. 3B, Fig. S13-S14**). Upon differentiating UB organoids to CD organoids, UPC-derived STR organoids expanded to form fluid-filled, cyst-like spheres. By comparison, static controls exhibited no gross morphological changes and EB-derived STR organoids appeared darker (**Fig. 3B, Fig. S13B**). The lack of change observed in static embedded organoids may suggest stabilization or confinement by the 3D matrix.

**Figure 3.**
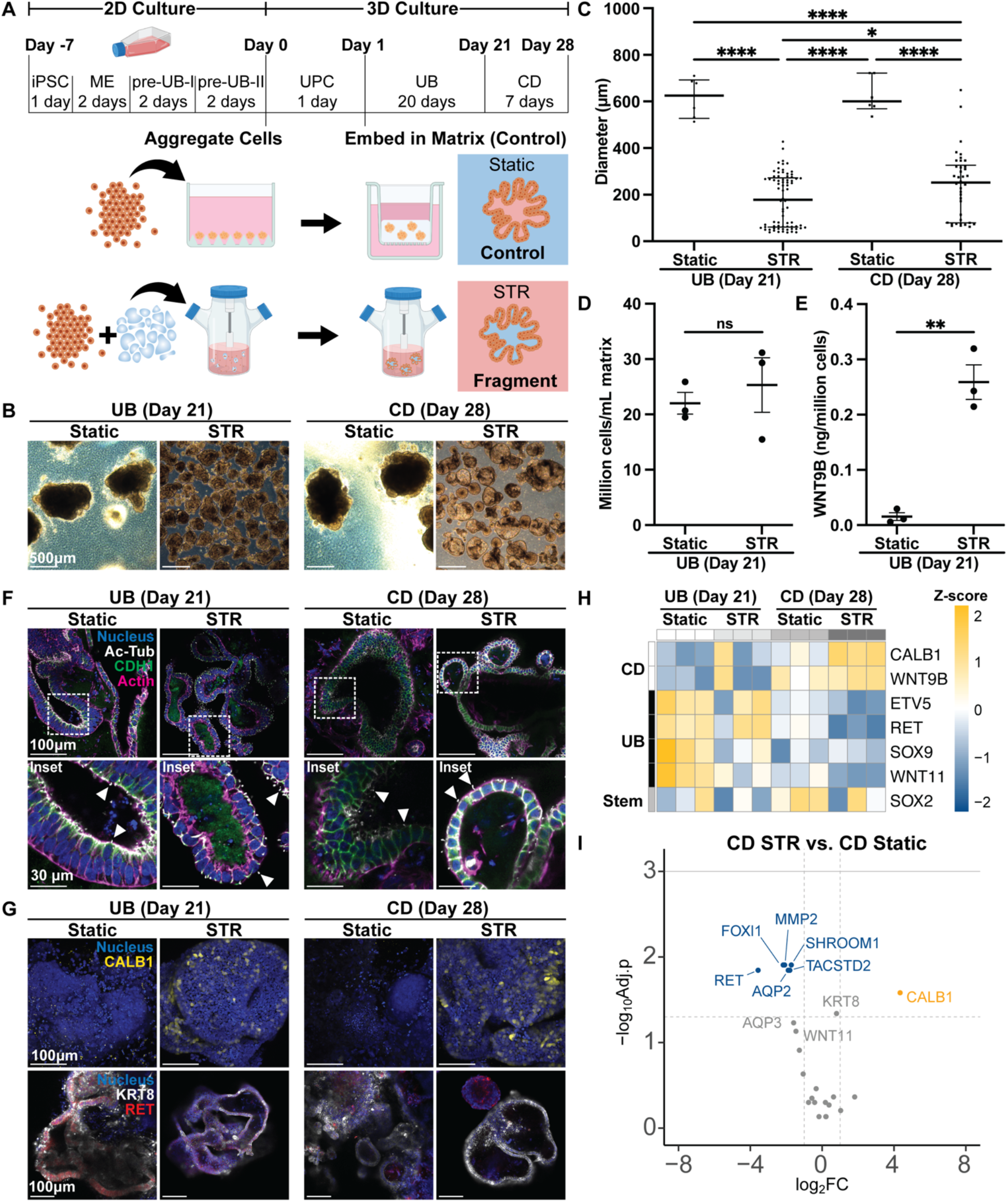
UB and CD organoid differentiation via matrix microcarriers in stirred bioreactors. A) Overview of culture workflow. UPCs are combined with fragmented matrix in an STR flask, cultured to day 21 in UB medium, then cultured to day 28 in CD medium. B) Morphology of UB and CD organoids (STR and static control) after 21 days in culture and CD organoids after an additional 7 days in culture. Scale = 500 μm. C) Diameter of organoids in each condition. Bars represent median and interquartile range. *p<0.05, p****<0.0001 using two-way ANOVA followed by Sidak’s multiple comparisons test. N = 3 independent differentiations. D) Cell yield normalized to volume of BsM used during culture for embedding organoids or as matrix microcarriers. Bars represent mean and SEM. No statistical significance using Welch’s t-test. N = 3 independent differentiations. E) WNT9B concentration in medium normalized to cell density in each condition measured by ELISA. Bars represent mean and SEM. p**<0.01 using Welch’s t-test. N = 3 independent differentiations. F) Representative immunofluorescence images of organoids in control and STR conditions. White box indicates inset. Blue = nucleus, white = Ac-Tub, green = CDH1, magenta = actin. Images are single slices. Scale = 100 μm. Inset shows apical surfaces oriented internally or externally. White arrows mark single primary cilium and apical surface. Scale = 30 μm. G) Representative immunofluorescence images of organoids in control and STR conditions. Blue = nucleus, yellow = CALB1, white = KRT8, red = RET. Images are maximum intensity projections (CALB1) or single slices (KRT8/RET). Scale = 100 μm. H) Heatmap of key markers for UB, CD, and stemness (stem) populations in each culture condition. N = 3 independent differentiations. Colors correspond to up-(orange) and down-(blue) regulated genes scaled to the standardized *Z* score. I) Gene expression in CD organoids cultured on fragment microcarriers in STRs relative to static embedded controls. Only UB and CD genes shown. Horizontal dashed line marks adj. p<0.05, horizontal solid line marks adj. p<0.01, vertical dashed line marks 2-fold change. Data points represent individual genes from N = 3 independent experiments, where *p* values are adjusted by the Benjamini-Hochberg correction.

To further characterize STR organoids, we quantified UB organoid size, yield, and growth factor secretion (**Fig. 3C-E, Fig. S13C-E**). Both STR UPC- and EB-derived organoids are significantly smaller than control (static embedded) organoids (**Fig. 3B-C, Fig. S13B-C, Fig. S14**), which may be useful given challenges associated with oxygen/nutrient diffusion in larger organoids. UPC-derived STR organoids produced similar cell yields as static embedded organoids when normalized by the total volume of matrix used in culture (**Fig. 3D, Fig. S13D**). While not statistically significant, organoids derived from EBs contained fewer cells per starting matrix volume, suggesting that this method may yield overall fewer differentiated cells (**Fig. S13D**). Next, we investigated WNT9B secretion to assess whether STR UB organoids exhibit functional differences compared to static embedded organoids. WNT9B is a marker of UB stalk populations and a powerful driver of nephrogenesis.^41^ When controlled for cell density, both UPC- and EB-derived STR UB organoids secreted more WNT9B than static embedded organoids (**Fig. 3E, Fig. S13E**), which fell below the assay detection limit. Hence, STR organoids derived from UPCs showed enhanced maturation and comparable yields to static controls. By contrast, STR organoids derived from EBs exhibited enhanced maturation, yet lower yields than those derived from UPCs.

Immunofluorescence staining revealed that while all organoids expressed key markers of UB/CD epithelium such as CDH1^+^ cell-cell contacts and GATA3^+^ nuclei at the protein level, their morphology and expression of key markers varied (**Fig. 3F-G, Figs. S15-S16**). For example, orientation of primary cilia (marked by acetylated tubulin, Ac-Tub) is generally inward for embedded organoids and outward for STR organoids, indicative of a reversal in their apical-basolateral polarization (**Fig. 3F, Fig. S15A**). We posit that changes in UB/CD cell orientation likely arise due to basolateral mechanical and adhesive cues present in the BsM. Consistent with transcriptomic data, STR CD organoids show an upregulation of CALB1 relative to static controls, while RET expression is generally higher in UB organoids compared to CD organoids (**Fig. 3G-H, Fig. S15B, Figs. S17-S18**). In contrast, expression of KRT8 and AQP2 was qualitatively similar in all conditions (**Fig. 3G, Fig. S15B, Fig. S16B**).

Transcriptomic analysis further revealed that CD organoids exhibit upregulation of stalk markers (*e.g*., *CALB1, WNT9B*) and downregulation of tip markers (*e.g., RET, SOX9, WNT11*) relative to UB organoids under all conditions (**Fig. 3H, Figs. S17-S18**). Moreover, CD organoids derived from UPCs cultured on microcarriers in suspension exhibited upregulation of *CALB1* and downregulation of *RET* compared to static embedded CD organoids (**Fig. 3I, Figs. S17-S18**). Interestingly, *AQP2* was also downregulated (**Fig. 3I**). If the morphology observed by phase contrast is indeed a result of increased fluid transport, changes in *AQP2* expression may be a compensatory effect to modulate fluid transport. These results were generally consistent with the previously observed trend that STR culture promotes expression of maturation markers, but that other upstream cues (*e.g*., transport) may obscure the effect. Generally, these findings indicate that differentiation of CD organoids in stirred bioreactors increases their maturation consistent with trends observed for UB organoids.

## Discussion

We have addressed a critical need by creating scalable UB/CD organoid differentiation protocols. By combining UPCs (or EBs) with fragmented BsM microcarriers within STRs, we have generated UB/CD organoids with enhanced maturation, as evidenced by changes in marker expression at both the gene and protein level. BsM microcarriers not only promote more homogenous expression of key UB markers and epithelial morphology but also enable control over apical-basolateral polarization. While all protocols tested give rise to epithelial populations that express key UB/CD markers (*e.g*., KRT8, CDH1, GATA3), microcarrier-based STR organoids exhibited enhanced expression of maturation and stalk phenotype markers (*e.g*., CALB1, WNT9B). Importantly, these organoids contained lumenized tubular networks that are contiguous with the outer monolayer, suggesting direct exposure of apical lumens to culture medium. This apical-out orientation, observed when using fragmented microcarriers in STR culture, may enable cells to sense fluidic shear stress via primary cilium. It is known that apical shear stress and luminal stretch can both increase intracellular calcium, leading to increases in potassium secretion^42,43^, and mechanically induced intracellular peak [Ca^2+^] increases during development.^44,45^ Luminal flow is a central driving force of tubule elongation during branching morphogenesis in developing kidneys *in vivo*.^46^

The pronounced effects of microenvironment in STR culture on UB/CD organoid differentiation raises an important question, namely: what are the specific roles of fluid flow, shear stress, and nutrient supply/oxygenation in cellular responses during differentiation in stirred bioreactors? To unravel these complex effects, one could leverage UB/CD organoid-on-chip models to systematically vary fluid flow rate, shear stress, and media composition. Another emerging question is how generalizable is this fragmented microcarrier-STR approach to other hiPSC lines and epithelial-based differentiation protocols (*e.g.*, lung or intestinal organoids)? It is anticipated that our approach could be adopted widely, albeit with specific matrix compositions developed for different organoids of interest. Finally, we speculate that designing microcarriers with controlled geometry, adhesion, and mechanics may allow one to further modulate the cellular microenvironment during STR differentiation enabling the scalable production of designer epithelial organoids.

In summary, we have established a robust and scalable method for generating UB/CD organoids by combining hiPSCs with fragmented BsM microcarriers in stirred bioreactors. This method results in kidney organoids with improved morphological development and maturation, which may find potential use in drug discovery, disease modeling, and therapeutic repair.

## Materials and methods

An overview of all utilized reagents, resources, and antibodies can be found in **Tables S3-S4**.

### Growth factor preparation

The following growth factors were reconstituted in DPBS^+/+^ containing 0.1% w/v BSA: 1 mg/mL FGF2, 100 μg/mL FGF7, 100 μg/mL GDNF, 100 μg/mL Rspo1, and 200 μg/mL EGF. The following small molecules were reconstituted in DMSO: 10 mM Y-27632, 1 mM LDN-193189, 10 mM CHIR-99021, 10 mM TTNPB, 20 mM A83-01, 10 mM JAK inhibitor I, 20 mM SB202190, and 10mM aldosterone.

### iPSC culture

BJFF.6 human iPSCs (male, provided by Prof. Sanjay Jain at Washington University) were maintained on plates coated with 1% Matrigel in DMEM/F12 and cultured in mTeSR Plus medium. For regular cell maintenance, cells were passaged at 60-80% confluency with ReLeSR according to manufacturer’s directions and split at ratios from 1:8 to 1:20. Frozen stocks were maintained in a 1:1 cell embryonic freezing medium and mTeSR Plus medium with 10 μM Y-27632. Normal karyotype of cells was validated by WiCell. Mycoplasma screening was performed at least every six months (Mycoalert, Lonza). Cells were passaged for no more than 15 times after most recent karyotype analysis.

### Generation of UB organoids in 96-well plates and cell culture inserts

iPSCs were differentiated towards UB organoids following the protocol of Zeng & Huang *et al.* with adaptations as necessary for STR culture.^21^ Briefly, iPSCs were washed with DPBS without Ca^2+^ and Mg^2+^ (DPBS^-/-^) and dissociated from the flask with Accumax solution for 5-10 min at 37 °C, 5% CO_2_. Cells were gently collected and an equal volume of mTeSR Plus medium with 10 µm Y-27632 was added. Cells were spun down at 200 g for 2 min and resuspended in mTeSR Plus with 10 µm Y-27632. Subsequently, 60,000 cells per well were seeded into a 1% Matrigel pre-coated 12-well plate, or 1.2 million cells into a T75 coated flask. Medium was replaced with pre-warmed mesendoderm (ME) medium the day after seeding (day -6). At day -4, the ME medium was replaced by pre-warmed pre-UB-I medium, and medium was refreshed the following day. At day -2, pre-UB-I medium was replaced with pre-UB-II medium, and medium was refreshed the following day. All media formulations can be found in **Table S1**. On day 0, differentiated cells were washed with DPBS^-/-^ and then dissociated with Accumax solution for 3-10 min at 37 °C. At this point, cells are either sorted via FACS or directly aggregated (**Fig. S6**). For cell sorting, cold FACS buffer (DPBS^+/+^ with 1% P/S and 2% FBS) was added in equal volume to the collected cells and cells were spun down at 300 g for 5 min at 4 °C. The cells were then resuspended in 2 mL of cold FACS buffer and 2 μL per initial 12 well plate of anti-CD117 PE-antibody was added. Samples and antibody were kept on ice for 30 min. The tube was gently tapped every 10 min to ensure mixing. Subsequently, samples were washed with cold FACS buffer and resuspended in 2 ml per plate cold FACS buffer supplemented with 1:4000 DAPI. Cells were passed through a 40 µm cell strainer and transferred into a FACS tube. CD117^+^DAPI^-^ cells were sorted with either a Beckman Coulter MoFlo Astrios EQ or a BD FACS Aria in the Bauer Core facility and collected in FACS buffer. Cells were spun down at 300 g for 5 min and resuspended in UBCM (**Table S1**) with 10 μM Y-27632 (UBCM-Y). Cells were counted using an automated cell counter (Countess 3 FL, Thermo Fisher Scientific) and aggregated in 96-well ultra-low adhesion plates or AggreWell800 microwell plates at 4,000 cells per aggregate (**Fig. 1A**) per manufacturer’s protocol. On day 1, UPC aggregates in 96-wells were individually embedded in 30 μL of 50% Matrigel diluted in DMEM/F12, HEPES, incubated for 30 min at 37 °C, 5% CO_2_, then cultured in 200 μL UBCM. Microwell-derived aggregates were cultured on PET cell culture inserts in 6-well plates by first creating a 1 mL base layer of 50% Matrigel on the insert and incubating for 30 min at 37 °C, 5% CO_2_. Approximately 300 aggregates were resuspended in 500 μL 50% Matrigel and transferred to the gel base layer, incubated for 30 min at 37 °C, 5% CO_2_, then cultured basolaterally with 2-2.5 mL of UBCM. An important limitation of the protocol thus far was the need to sort CD117^+^ UPCs at day 0, which typically constituted 50-80% of cells. We first compared organoids derived from CD117^+^ cells to CD117^-^ sorted cells and saw that the CD117^-^ cells still gave rise to organoids with buds with lumens observable by phase imaging. However, organoids generated from CD117^-^ cells were smaller than those from CD117^+^ cells (**Fig. S6A**). When comparing organoids derived from CD117^+^ cells to unsorted cells, no qualitative differences in organoid size or morphology were observed (**Fig. S6B**). After 21 days in culture, transcriptomic analysis of organoids derived from CD117^+^ sorted cells and unsorted cells revealed no significant differences in gene expression (**Fig. S6C**). Together, these results suggest that CD117^+^ sorting is not necessary when differentiations are robust (>50% CD117^+^ at day 0). We eliminated the FACS sorting step from all subsequent experiments.

### Generation of UB/CD organoids in shaker and STR cultures from UPCs

UPCs were generated as described above. On day 0, UPCs were aggregated in AggreWell800 microwells (4,000 cells/microwell) or non-adherent T25 flasks on an orbital shaker (VWR 3500 Advanced Digital Shaker) with or without matrix microcarrier fragments (**Fig. 2A**). Cells were resuspended at 0.5-1.0 million cells/mL in 12 mL UBCM-Y and cultured for 24 h at 60-70 RPM to generate cell aggregates (**Fig. S8**). In parallel, cells were resuspended at ∼1 million cells/mL in 11.5 mL UBCM-Y with 500 μL matrix fragments composed of Geltrex and cultured for 24 h at 60 RPM, 37 °C, 5% CO_2_ to promote cell adhesion to the matrix. On day 1, microwell-derived aggregates were either encapsulated in 100% Geltrex droplets, left static in microwells, or embedded in 50% Geltrex on cell culture inserts as a control. To generate droplets, individual aggregates were suspended in ∼2 uL Geltrex on a polystyrene surface, inverted and incubated at 37 °C, 5% CO_2_ for 1 h, then transferred using a cell scraper into a non-adherent T25 flask containing 12 mL UBCM (**Table S1, Fig. S7**). Shaker-derived aggregates (with and without microcarriers) were centrifuged at 100 g for 3 min to remove the culture medium, then resuspended in 12 mL UBCM. Shaker cultures, cell culture inserts, and static microwells were refreshed every 2-3 days with 12, 2.5, and 2 mL UBCM, respectively. On day 21, UB organoids were fixed and lysed for immunostaining and transcriptomic analysis, respectively. The microcarrier method was translated to STRs by scaling the volume of medium and matrix to 60 mL and 2.5 mL, respectively, with a cell seeding density of 1 million cells/mL in UBCM-Y at 60 RPM, 37 °C, 5% CO_2_ (**Fig. 3A**). After 24 h, UBCM-Y was replaced with UBCM and UB organoids were refreshed with 60 mL UBCM every 2-3 days until day 21, then collected for analysis or further differentiated to CD organoids for 7 days. CD differentiation was performed as described in Zeng & Huang *et al*.^21^ On day 28, CD organoids were collected for immunostaining and transcriptomic analysis.

### Generation of UB/CD organoids in shaker and STR cultures from EBs

hiPSCs were differentiated to UB organoids by adapting the above protocol to an entirely 3D culture system using EBs. On day -7, iPSCs were washed with DPBS^-/-^ and dissociated from the flask with Accumax solution for 5-10 min at 37 °C, 5% CO_2_. Cells were gently collected and an equal volume of mTeSR Plus medium with 10 µm Y-27632 was added. Cells were centrifuged at 200 g for 2 min and resuspended in mTeSR Plus with 10 µm Y-27632. Cells were seeded into non-adherent T25 flasks on an orbital shaker at 1 million cells/mL in either 12 mL medium alone or 11.5 mL medium with 500 μL Geltrex microcarriers and cultured overnight at 60 RPM, 37 °C, 5% CO_2_ (**Fig. S11A**). On day -6, the cultures were centrifuged at 100 g for 3 min to pellet the aggregates and microcarriers, and the medium was replaced with 12 mL pre-warmed ME medium. On day -4, the cultures were centrifuged and the ME medium was replaced with 12 mL of pre-warmed pre-UB-I medium, and medium was refreshed the following day. On day -2, the cultures were centrifuged and the pre-UB-I medium was replaced with 12 mL of pre-UB-II medium and refreshed the following day. On day 0, the cultures were centrifuged and the pre-UB-II medium was replaced with UBCM. Cultures were refreshed every 2-3 days with 12 mL UBCM. On day 21, UB organoids were fixed and lysed for immunostaining and transcriptomic analysis, respectively. To translate the microcarrier method to STR culture, the iPSC seeding density on day -7 was reduced to 50,000 cells/mL to account for proliferation during differentiation, and the volume of medium and matrix was scaled to 60 mL and 2.5 mL, respectively (**Fig. S13A**). STRs were seeded at 50,000 cells/mL in UBCM-Y at 60 RPM, 37 °C, 5% CO_2_. After 24 h, UBCM-Y was replaced with UBCM and UB organoids were refreshed with 60 mL UBCM every 2-3 days until day 21, then collected for analysis or further differentiated to CD organoids for 7 days. On day 28, CD organoids were collected for immunostaining and transcriptomic analysis.

### Fragment preparation

Extracellular matrix microcarriers were generated by extrusion fragmentation (**Fig. S7**), as previously adapted from a protocol by Muir *et al.*^40,47^ for generating granulated bulk hydrogels. Pure Geltrex was transferred to a 5 mL syringe on ice, then incubated at 37 °C for 30 min to allow for cross-linking. The solid matrix was then extruded through a 32 G / 0.10 mm I.D. nozzle at ∼1 drop/second into the culture vessel containing UBCM-Y. Dissociated UPCs or hiPSCs were then combined with microcarriers in suspension, pipetted repeatedly to ensure even mixing of cells and separation of fragments, and cultured at 37°C, 5% CO2 on an orbital shaker or STR stirring plate.

### Organoid area analysis

Phase contrast images of organoids were acquired on days 1 (EB), 21 (UB), and 28 (CD) using a 4X objective (Leica DM IL LED Inverted Laboratory Microscope). Raw image files were converted from .lif to .tif format in ImageJ, then imported into ilastik-1.4.0 for object classification. A training set was compiled using an image from each timepoint to train a pixel classification algorithm by manually identifying foreground (organoids) and background. The trained algorithm was used to classify objects in subsequent images, yielding a colorized mask overlay image containing only foreground and background. These outputs were then imported into ImageJ, converted to 8-bit binary images, followed by watershed analysis to separate neighboring organoids. ImageJ Particle Analysis was used to identify and segment organoids, and the Measurement tool was used to quantify 2D area of each organoid. Measurements were imported into GraphPad Prism 10, and 2D area was converted to diameter for plotting and statistical analysis. A threshold of 50 μm was applied to exclude dead cells and debris from quantitative analysis.

### Cell yield analysis

On days 21 and 28, the total STR culture was centrifuged at 100 g for 3 min and the supernatant was removed. The remaining pellet was centrifuged again at 200 g for 1 min, and the remaining supernatant was removed. The pellet was slowly pipetted using a P1000 micropipette with a wide-bore tip to measure the total tissue volume. A sample of 20-50 μL was removed, washed with DBPS^-/-^, and treated with 10 mL Accumax on an orbital shaker at 60 RPM, 37 °C, 5% CO_2_ for 30-45 min until cells were fully dissociated. Two counts were obtained using the automated cell counter and extrapolated to the total volume of tissue to measure the total cell yield of the STR. Measurements were imported into GraphPad Prism 10 for plotting and statistical analysis.

### RNA isolation and Nanostring analysis

For each condition, ∼10 organoids were collected. Organoids were rinsed with DBPS^+/+^. Adherent 2D iPSCs were first rinsed with DPBS^-/-^, then treated with Accumax, and finally pelleted at 300 g for 5 min. Supernatant was removed from organoids and iPSCs, then RNA was isolated using the RNeasy Plus Mini Kit according to manufacturer’s instructions. RNA concentration was assessed using both the Nanodrop 1000 spectrophotometer (NanoDrop Products, Thermo Scientific, Wilmington, DE) and Qubit (Life Technologies). A custom kidney gene panel was designed by NanoString and produced by IDT (**Table S2**) using probes containing 35–50 bp each. RNA quantification was performed by NanoString nCounter Elements™ with appropriate reagents as per the manufacturer’s instructions, using 100 ng RNA per run. RNA was hybridized with probe pools, hybridization buffer, and TagSet reagents in a total volume of 30 μL and incubated at 67 °C for 20 h. Quantification data was analyzed by the nSolver software using a custom advanced analysis. No low-count data were omitted in the initial analysis. *ACTB*, *GAPDH*, and *TUBB* were marked as housekeeping genes. After processing the data in nSolver, low-count data was removed from further analysis. Low-count genes were defined as genes in which the mRNA count was below the threshold for at least one replicate in all experimental conditions. We considered genes as significantly differentially expressed when the adjusted *p* value was <0.05. Our study was powered at 0.8 to detect a 2-fold change in expression, with the number of biological replicates determined based on preliminary data. Data was plotted with R.

### Immunostaining

Prior to immunostaining, each sample was washed with DPBS^+/+^ and then fixed for 1 h with 10% buffered formalin solution. The fixative was removed by 3 washes in DPBS^+/+^ for ∼2 h each, and samples were then blocked and permeabilized for 24 h with 1% v/v donkey serum in DPBS^+/+^ with 0.125% v/v Triton X-100. Primary antibodies were incubated with the samples for 48 h at 4 °C in a staining solution (0.5% w/v BSA and 0.125% v/v Triton X-100 in DPBS^+/+^). Samples were washed 3 times in DPBS^+/+^ for ∼2 h each. Secondary antibodies were incubated with the samples for 48 h at 4 °C in staining solution. Samples were counterstained with DAPI and then washed 3 times in DPBS^+/+^ for ∼2 h each before imaging. Confocal images were acquired on a Zeiss LSM 710. Imaris was used to visualize 3D projections, stacks, and rendering. ImageJ was used for processing and quantification.

### Protein quantification

An aliquot of medium was collected from each STR flask at 21 days in culture, exactly 72 hours after receiving a full volume (62.5 mL) of fresh culture medium. These undiluted media were assayed via ELISA for WNT9B according to manufacturer’s guidelines using two technical replicates on undiluted media. Absorbance was measured and analyzed using spectrophotometry (BioTek, Synergy HTX Multimode Reader). Raw values were imported into GraphPad Prism 10, converted to WNT9B concentration (ng/mL) using the ELISA standard curve, extrapolated to the total medium volume (ng), then normalized to the total number of cells in the flask (ng/million cells). Normalized values were plotted and statistical analysis was performed using GraphPad Prism 10.

### Statistical analysis

The number of biological replicates necessary to achieve power of 0.8 were estimated using G*Power using previous or preliminary data. Statistical significance was calculated by the indicated tests using GraphPad Prism 10. For all experiments, N values correspond to the sum of replicates collected over at least 3 independent experiments. Statistical details (exact value of N and what N represents, statistical test used for comparison, definition of center, dispersion and precision measures, and definition of significance) can be found in figure legends.

## Resource availability

### Lead contact

Requests for further information, resources, or reagents can be directed to corresponding author, Jennifer A. Lewis.

### Materials availability

The BJFF.6 cell line used is available from the lead contact upon request.

### Data and code availability

All data supporting key findings of this study are available within this paper and the supplemental information. Raw data and analysis codes are available from the lead contact upon reasonable request.

## Acknowledgements

This work was supported by the NIH Re(Building) a Kidney Consortium (NIH UC2DK126023), Vannevar Bush Faculty Fellowship Program sponsored by the Basic Research Office of the Assistant Secretary of Defense for Research and Engineering through the Office of Naval Research Grant N00014-21-1-2958, the NIH F32 Ruth L. Kirschstein National Research Service Award for Individual Postdoctoral Fellows (KJW, DK131821), and Dutch Research Council Rubicon grant (RG, 019.201EN.005). The authors would like to thank staff at the Bauer Core at Harvard University for assistance with cell sorting.

## Author contributions

Conceptualization, K.J.W., J.E.R., R.C.v.G., and J.A.L.; data collection and analysis, K.J.W., J.E.R., R.C.v.G., K.S., A.N.K., P.N., and J.A.L.; methodology, K.J.W., J.E.R., R.C.v.G., K.S.; writing – original draft, K.J.W., J.E.R., R.C.v.G., and J.A.L.; writing, review & editing, all authors; funding acquisition, K.J.W., R.C.v.G., and J.A.L.

---

**Supplemental Figure 1.**
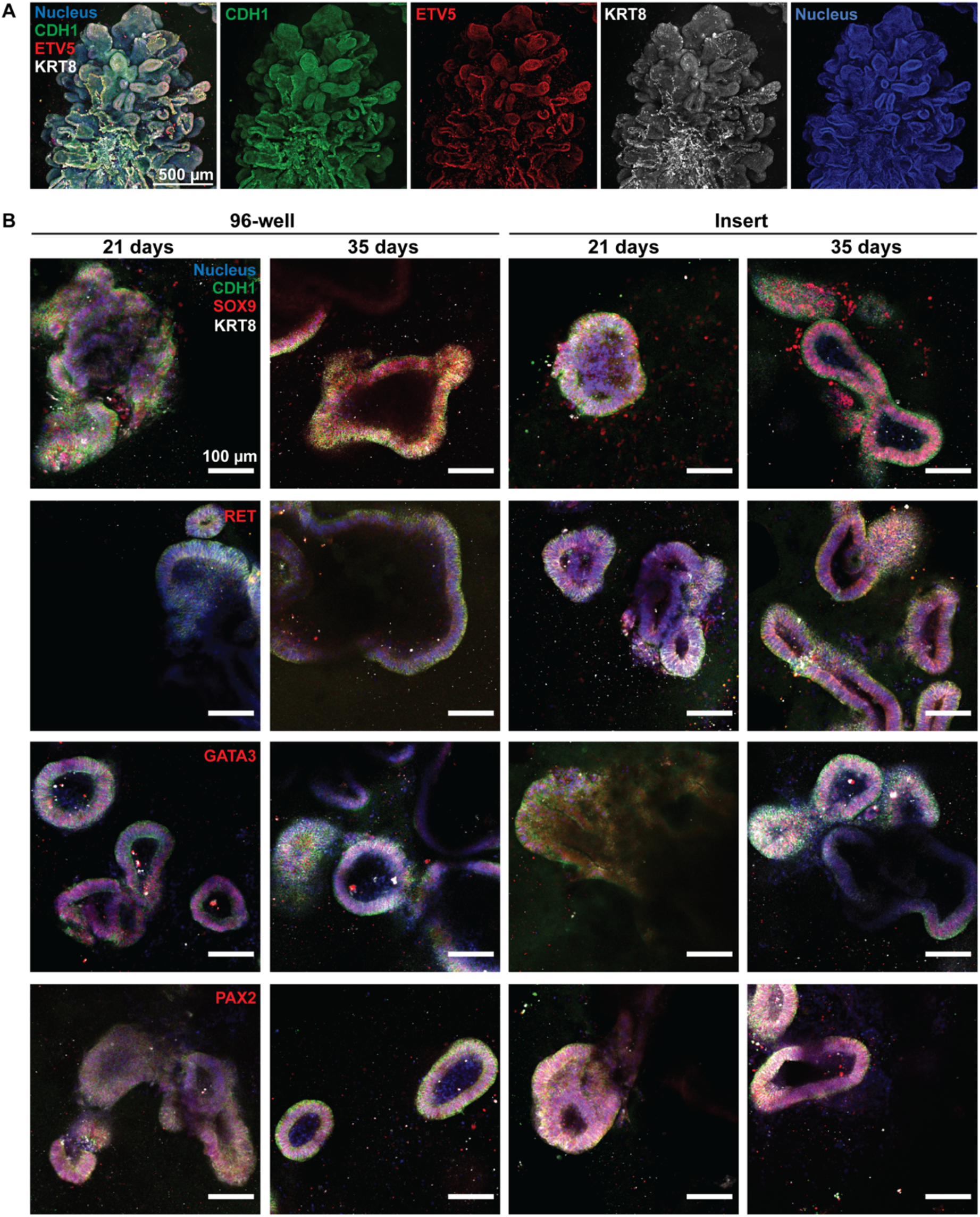
UB organoids embedded in matrix and cultured under static conditions. A) Immunofluorescence image (maximum intensity projection) of an embedded UB organoid cultured in an insert for 21 days. Green = CDH1, red = ETV5, white = KRT8, blue = nucleus. Scale = 500 μm. B) Immunofluorescence images (single slices) of UB organoids embedded in BsM and cultured under static conditions on 96-well plates and inserts for 21 days and 35 days. Blue = nucleus, green = CDH1, white = KRT8, red = SOX9, RET, GATA3, PAX2 (from top row to bottom row). Scale = 100 μm. N = 3 independent differentiations, all images shown from the same differentiation.

**Supplemental Figure 2.**
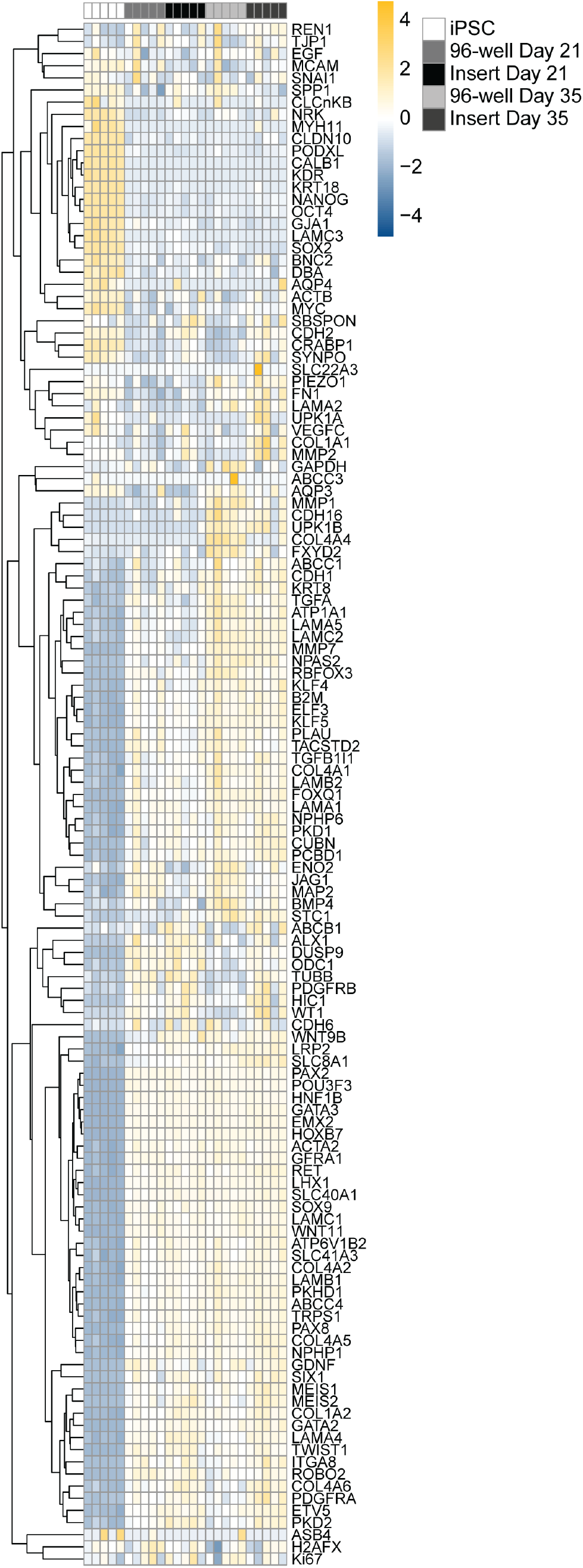
Transcriptomic analysis of UB organoids embedded in BsM and differentiated in 96-well plates and inserts at 21 and 35 days in culture compared to iPSCs. All tested genes above threshold are shown. N = 5 independent differentiations.

**Supplemental Figure 3.**
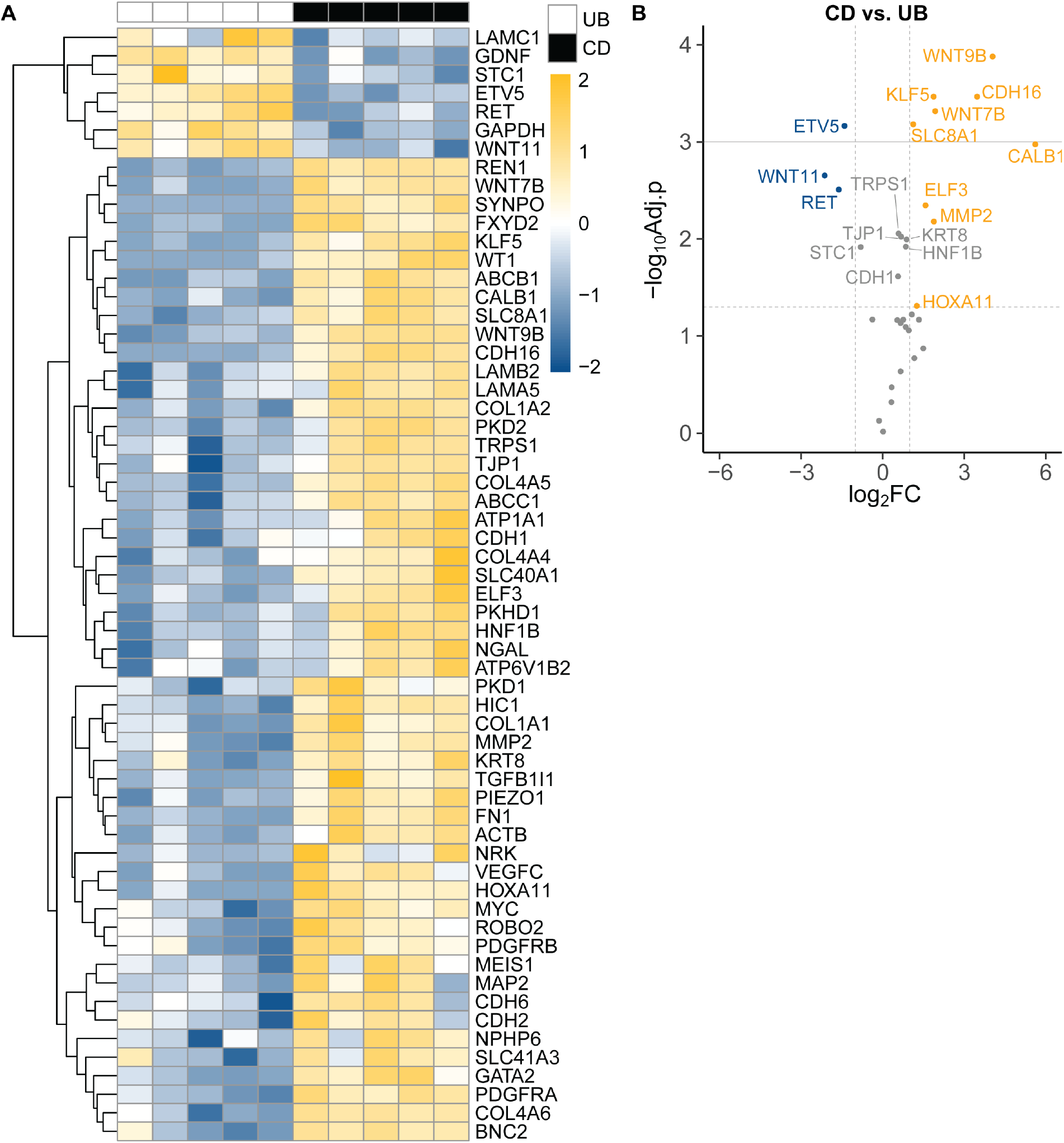
Transcriptomic analysis of static embedded UB and CD organoids at 28 days in culture. A) Transcriptomic analysis of all tested genes with expression above threshold. N = 5 independent differentiations. B) Differential gene expression of CD organoids relative to UB organoids. Only UB and CD genes shown. Horizontal dashed line marks adj. p<0.05, horizontal solid line marks adj. p<0.01, vertical dashed line marks 2-fold change. N = 5 independent differentiations.

**Supplemental Figure 4.**
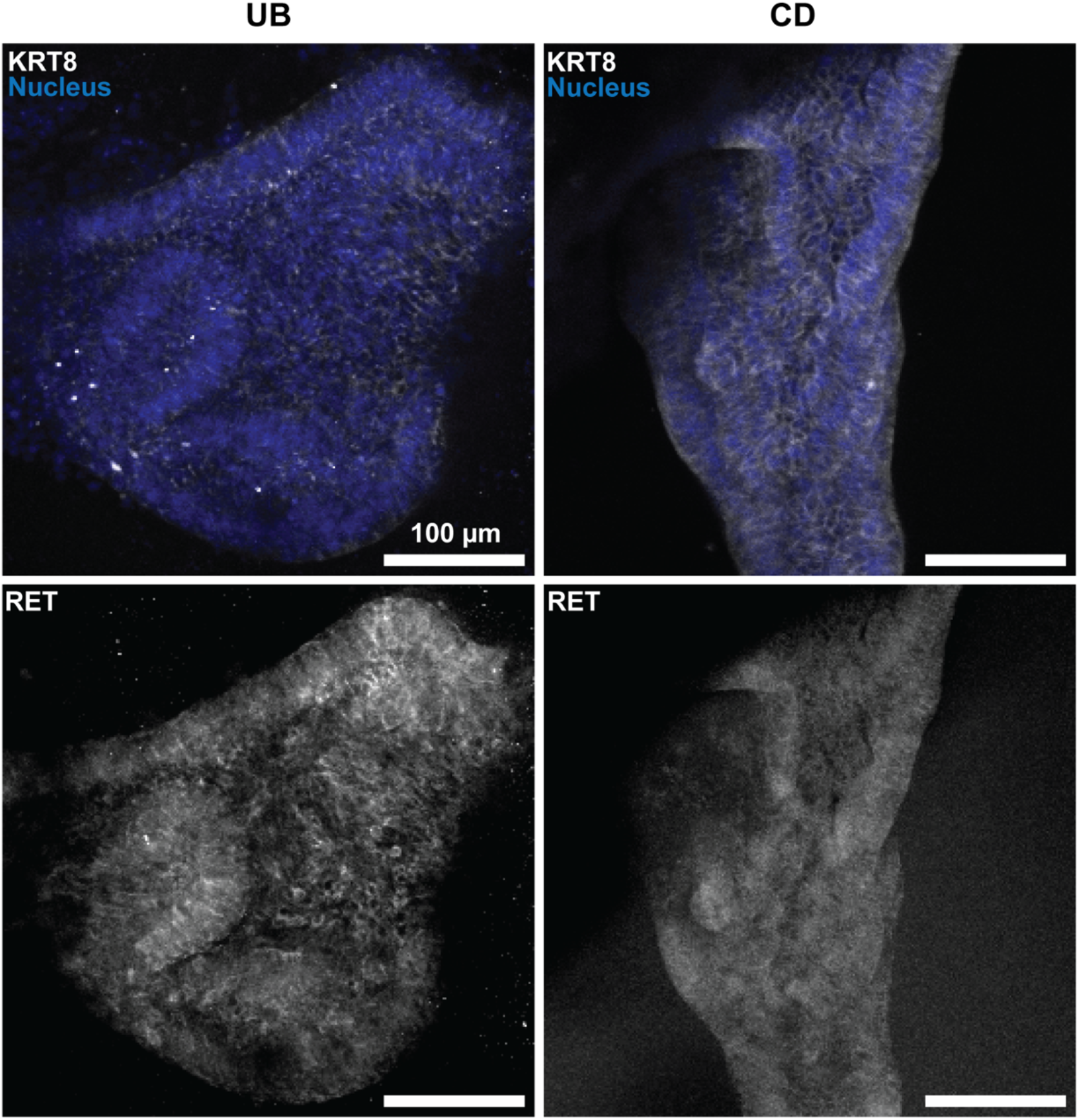
UB marker expression in UB and CD organoids. Immunofluorescence staining (maximum intensity projections) of UB stalk (KRT8) and UB tip (RET) markers in both UB and CD organoids. Blue = nucleus, white = KRT8, RET (from top row to bottom row). Scale = 100 μm. N = 3 independent differentiations, all images shown from the same differentiation.

**Supplemental Figure 5.**
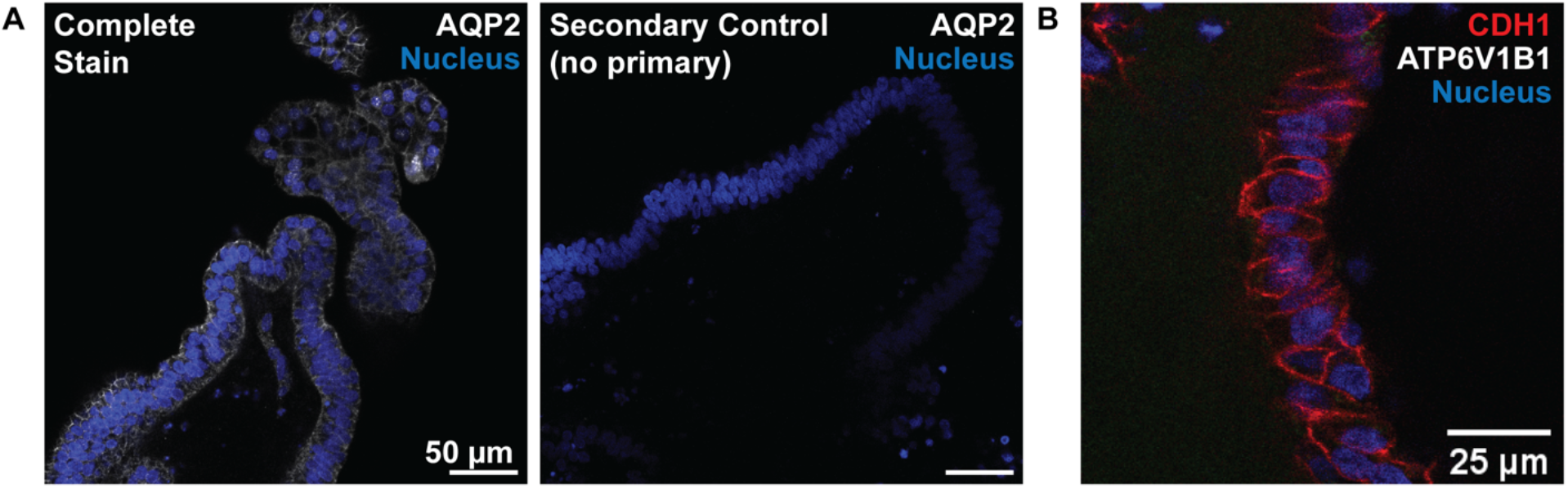
Characterization of principal-like cells in CD organoids. A) Immunofluorescence image of AQP2 expression in static organoids. Images showed with secondary control. White = AQP2, blue = nucleus. Scale = 50 μm. B) Absence of ATP6V1B1 in CD organoids. Red = CDH1, white = ATP6V1B1, blue = nucleus. Scale = 25 μm. N = 3 independent differentiations, all images shown from the same differentiation.

**Supplemental Figure 6.**
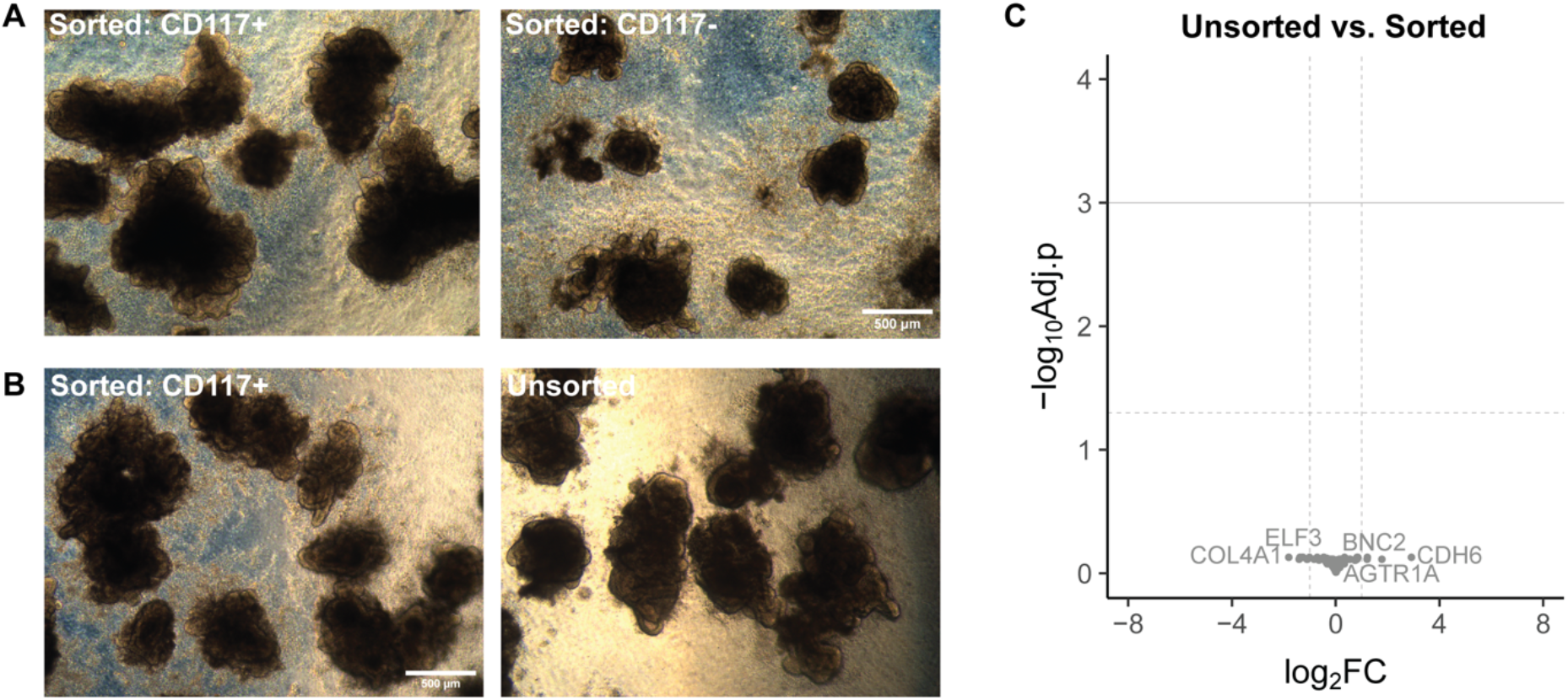
Comparison of CD117-sorted and unsorted UB organoids. A) Morphological comparison (brightfield images) of UB organoids at 21 days in culture derived from sorted cells that were CD117^+^ at day 0 compared to CD117^-^ at day 0. Scale = 500 μm. N = 3 independent differentiations, all images shown from the same differentiation. B) Morphological comparison of UB organoids at 21 days in culture derived from CD117^+^ cells at day 0 compared to unsorted cells. C) Gene expression in 21-day old organoids derived from unsorted cells at day 0 compared to sorted CD117^+^ cells at day 0. Horizontal dashed line marks adj. p<0.05, horizontal solid line marks adj. p<0.01, vertical dashed line marks 2-fold change. N = 3 independent differentiations.

**Supplemental Figure 7.**
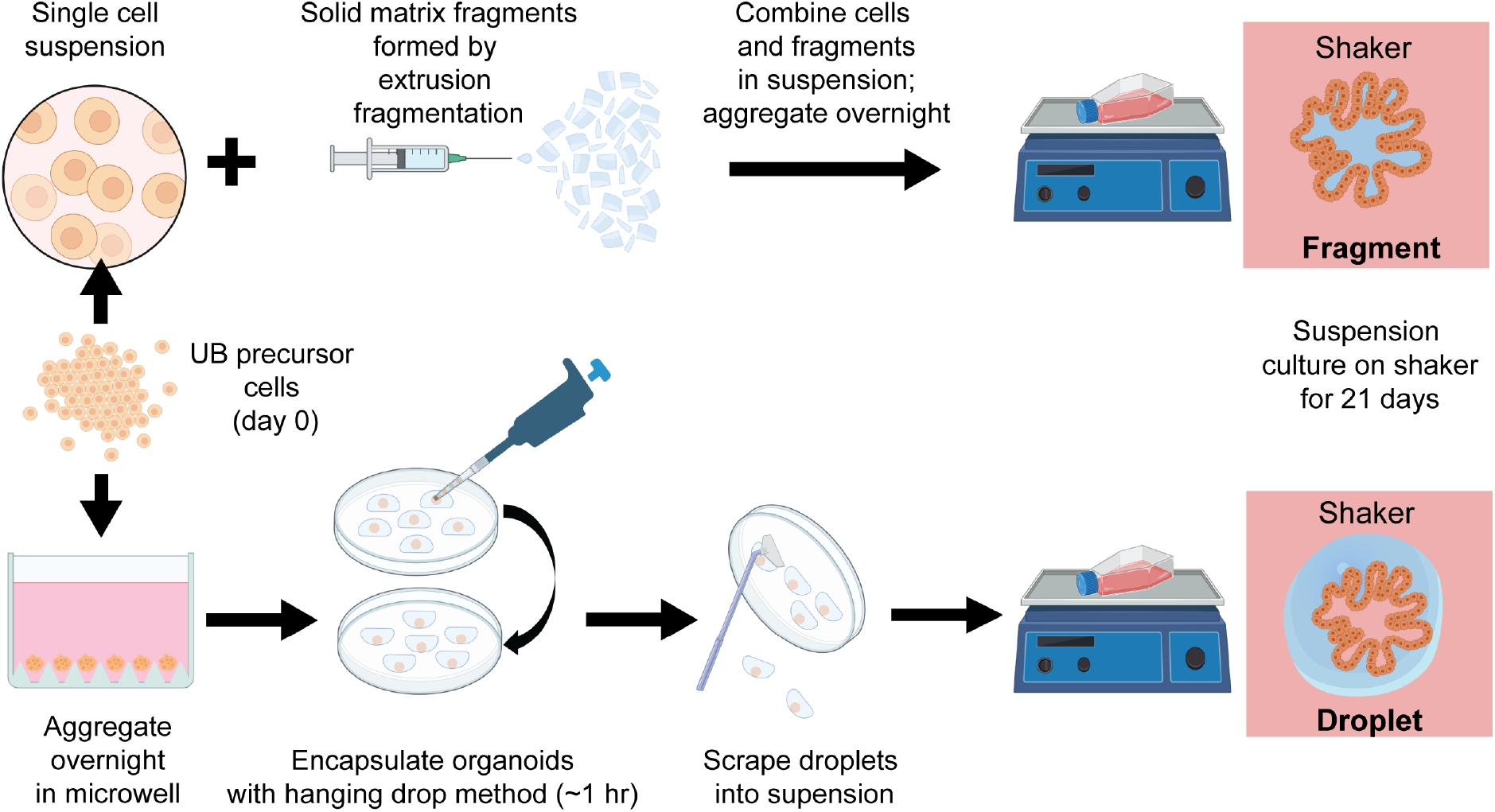
Matrix fragmentation and droplet encapsulation methods for UB/CD organoid differentiation.

**Supplemental Figure 8.**
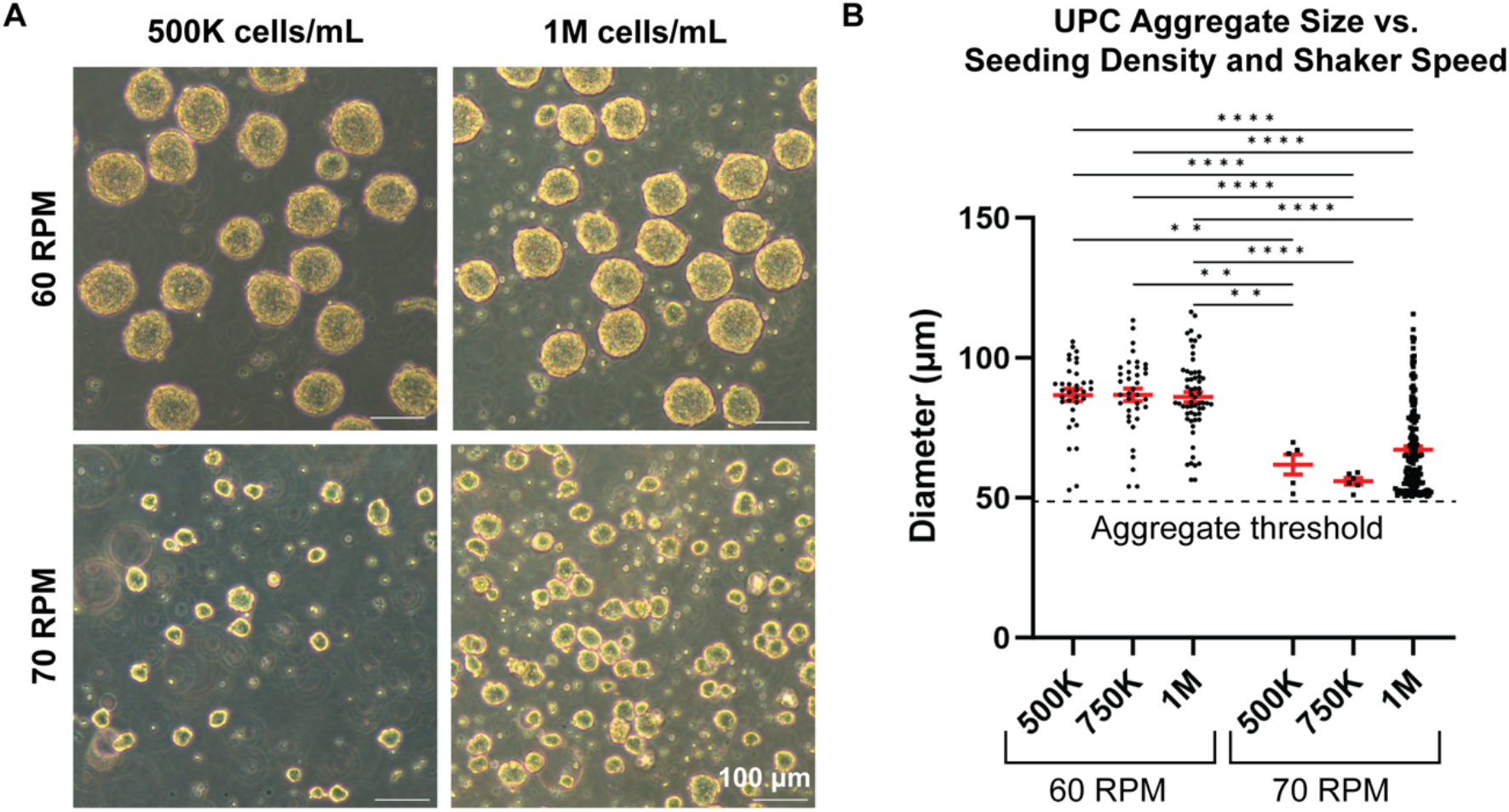
Aggregate formation in UPCs (day 0) without matrix. A) Aggregate morphology (brightfield images) with respect to seeding density and orbital shaker speed after overnight aggregation. N = 3 independent differentiations, all images shown from the same differentiation. Scale = 100 μm. B) Comparison of aggregate size with respect to initial cell seeding density and orbital shaker speed after overnight aggregation. Bars represent mean and SEM. **p<0.01, ****p<0.0001 using two-way ANOVA followed by Tukey’s multiple comparisons test. N ranged between 5 to 171 organoids per condition, which were collected across 3 independent differentiations.

**Supplemental Figure 9.**
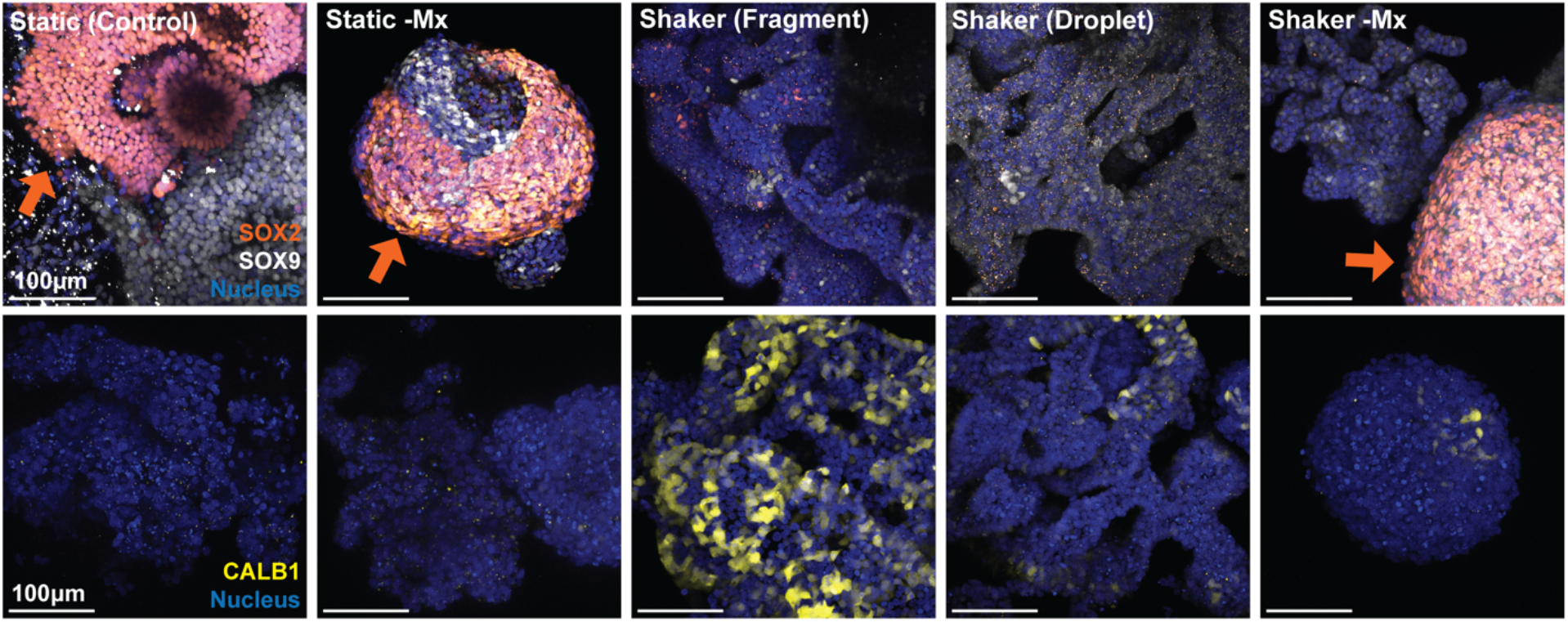
Adapting UB differentiation to suspension culture. Data is depicted from Figure 2, here including additional data on both static and shaker organoids cultured without matrix (-Mx) for clarity. Maximum intensity projections of key UB markers for organoids at 21 days in culture. Orange = SOX2, white = SOX9, yellow = CALB1, blue = nucleus. Orange arrows mark regions of SOX2^+^ cells. Scale = 100 μm. N = 3 independent differentiations, all images shown from the same differentiation.

**Supplemental Figure 10.**
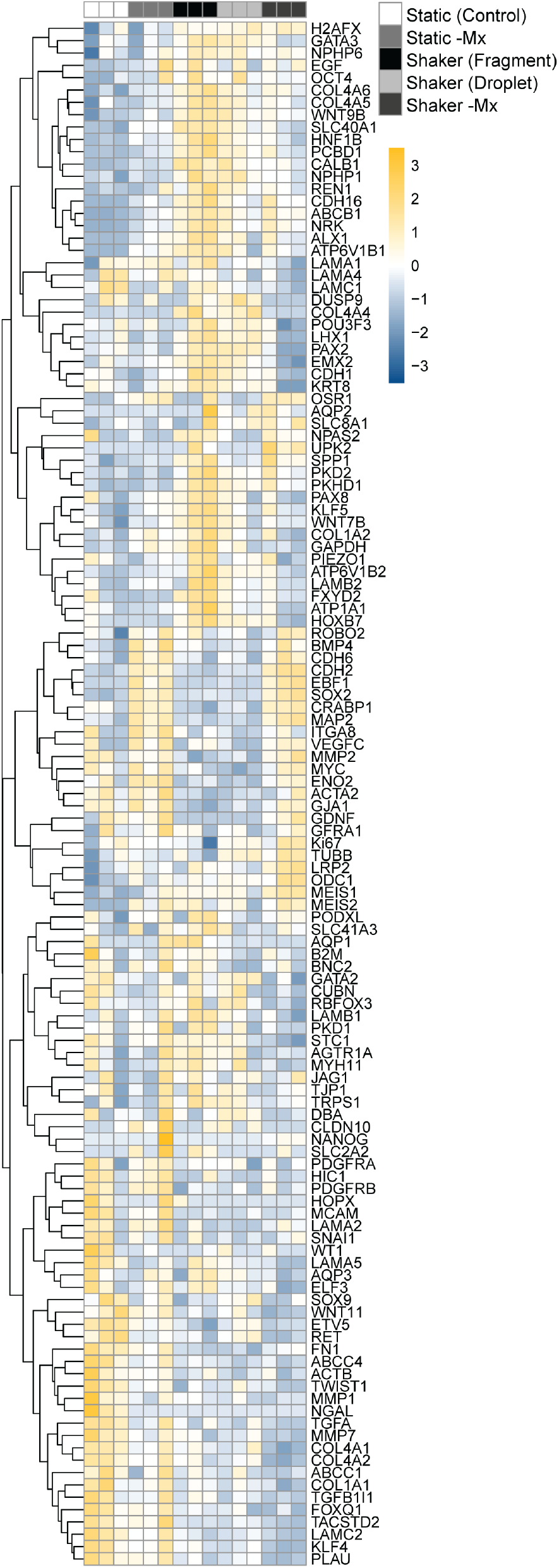
Transcriptomic analysis of UB organoids derived from UPCs cultured on orbital shakers with and without matrix microcarriers and static embedded controls. All tested genes above threshold are shown. N = 3 independent differentiations.

**Supplemental Figure 11.**
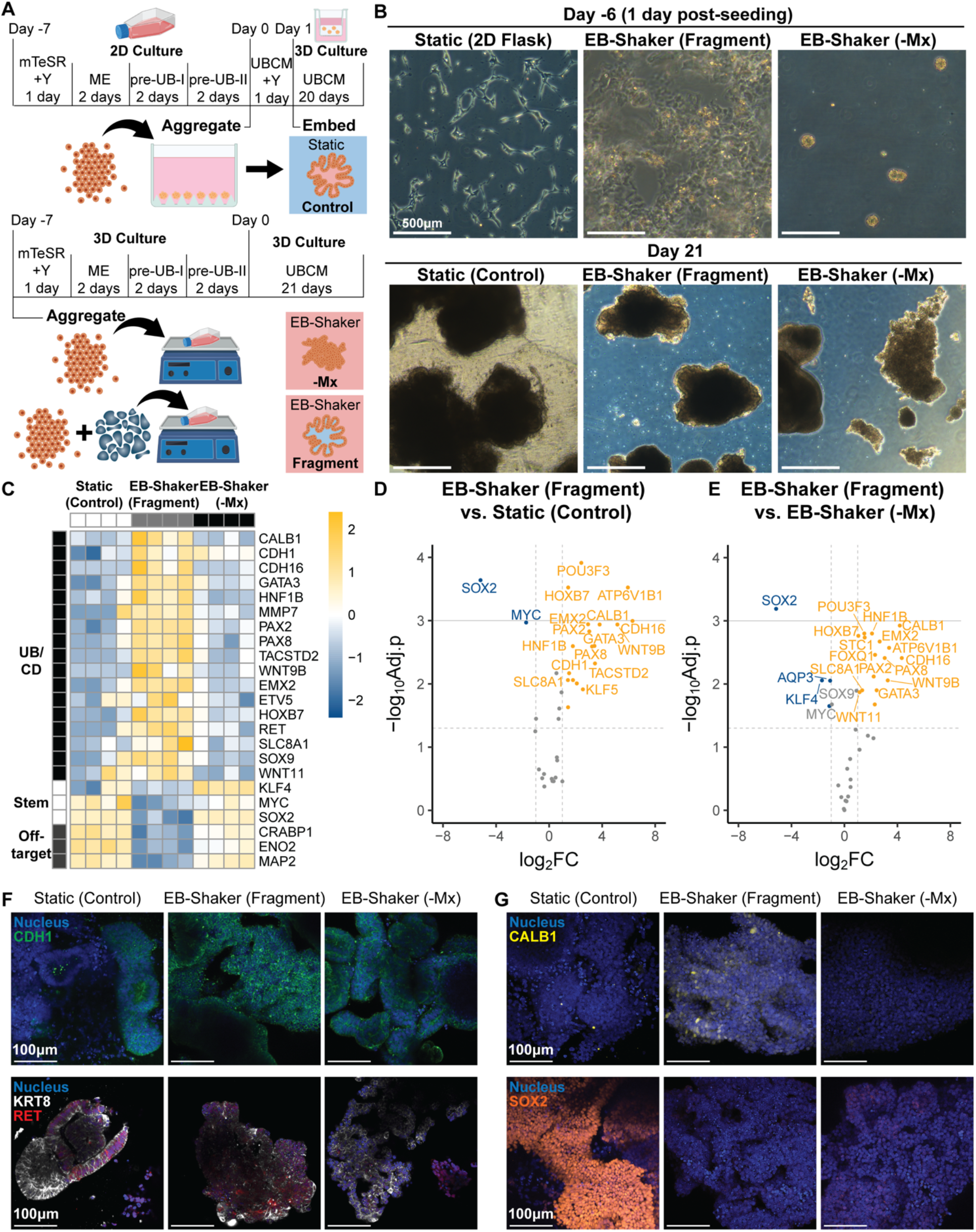
Adapting UB 2D differentiation and 3D embedded culture to suspension culture using matrix microcarriers. A) Overview of culture conditions. iPSCs were differentiated on 2D plastic and then embedded in static BsM in cell culture inserts served as a control. For shaker conditions, iPSCs (day -7 of protocol) were added to a culture flask with or without (-Mx) fragmented matrix and cultured on an orbital shaker. B) Morphology (brightfield images) of iPSCs 1 day after seeding (day -6) and UB organoids after 21 days in culture. While iPSCs in 2D flasks (control) form small colonies, iPSCs in shaker conditions form aggregates in the absence of matrix or adhere to matrix in the presence of fragments. By day 21, large, cellular organoids have formed. Scale = 500 μm. C) Transcriptomic analysis of key markers for UB, CD, stemness (stem), and off-target neuronal populations in each culture condition. N = 4 independent differentiations. D) Gene expression in embryoid-derived organoids cultured on fragment microcarriers on an orbital shaker relative to static embedded controls. Only UB and CD genes shown. E) Gene expression in embryoid-derived organoids cultured on an orbital shaker with fragment microcarriers relative to organoids without microcarriers (-Mx). For D and E, horizontal dashed line marks adj. p<0.05, horizontal solid line marks adj. p<0.01, vertical dashed line marks 2-fold change. N = 4 independent differentiations. F) Immunofluorescence images (single slices) of UB organoids in all conditions after 21 days in culture. Blue = nucleus, green = CDH1, white = KRT8, red = RET. Scale = 100 μm. G) Immunofluorescence images (maximum intensity projections) of UB organoids in all conditions after 21 days in culture. Blue = nucleus, yellow = CALB1, orange = SOX2. Scale = 100 μm. For F and G, N = 4 independent differentiations, all images shown from the same differentiation.

**Supplemental Figure 12.**
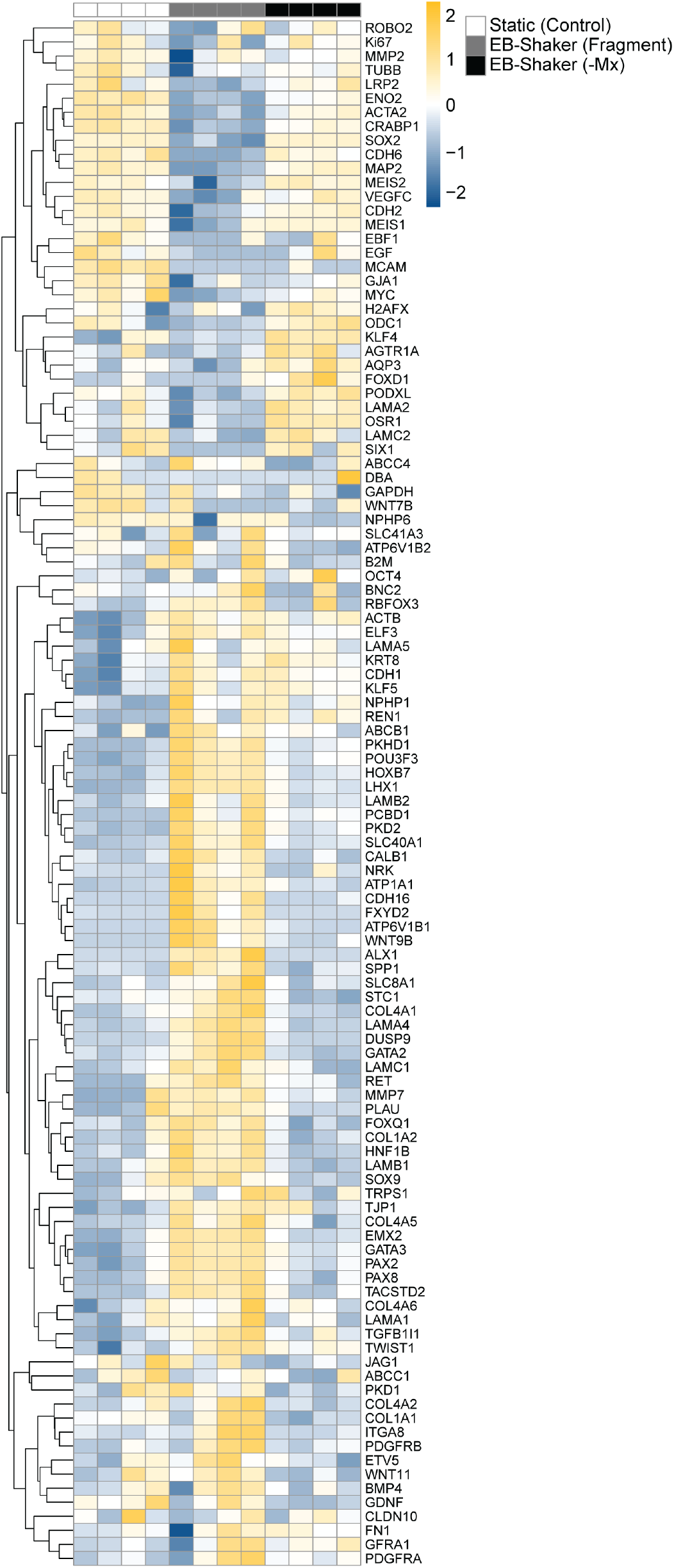
Transcriptomic analysis of UB organoids on orbital shakers derived from EBs and static embedded controls with and without (-Mx) matrix microcarriers. All tested genes above threshold are shown. N = 4 independent differentiations.

**Supplemental Figure 13.**
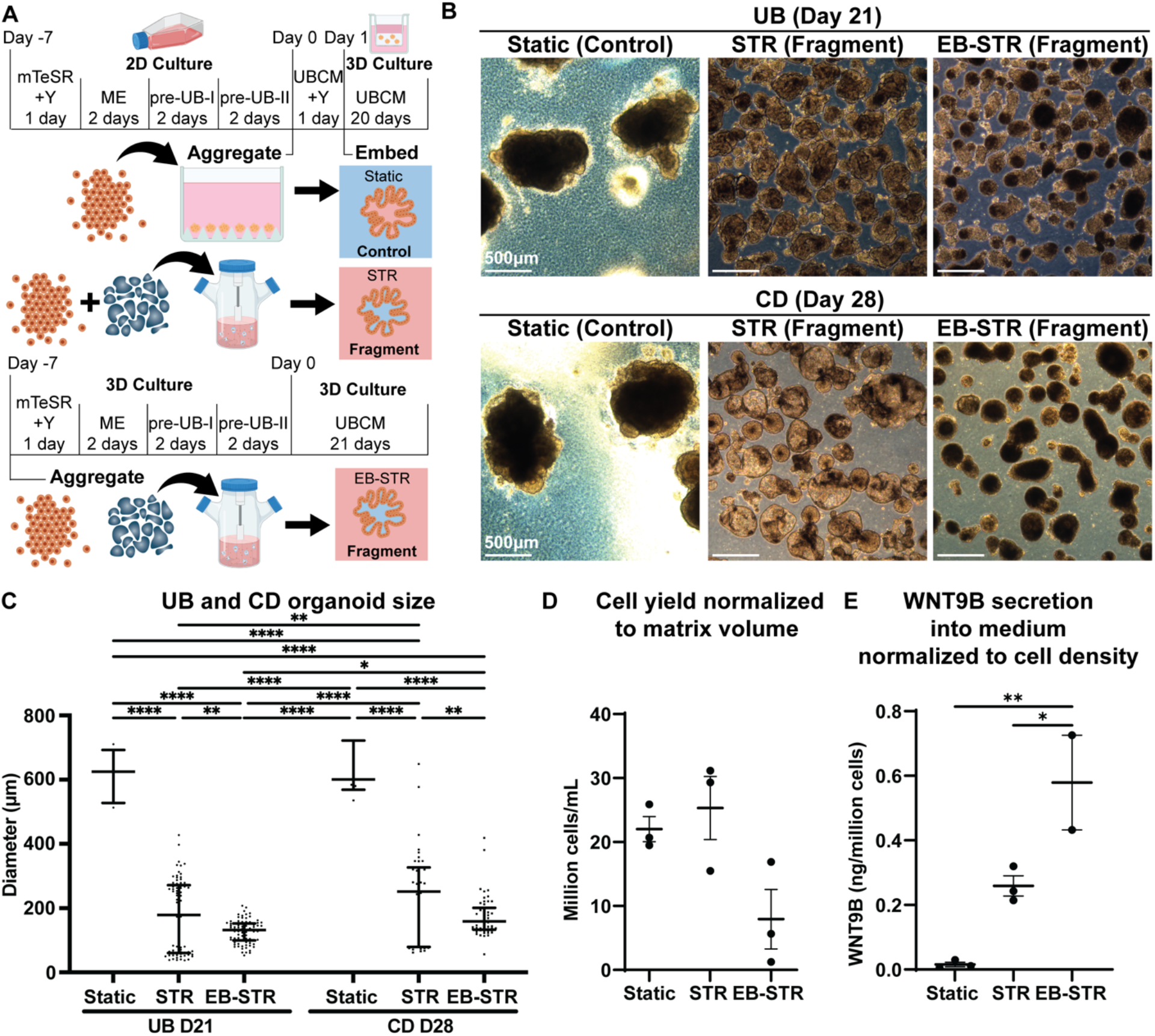
Scaling UB and CD organoid culture to STRs. Data is depicted from Figure 3, here including additional data on EB-STR (Fragment) for clarity. A) Overview of culture conditions: static embedded organoids in cell culture inserts compared to organoids cultured in STRs with fragment microcarriers starting with either UPCs (day 0) or EBs (day -7). At day 21, all organoids are cultured for an additional 7 days in collecting duct medium. B) Morphology (brightfield images) of UB and CD organoids after 21 and 28 days in culture, respectively. Scale = 500 μm. N = 3 independent differentiations, all images shown from the same differentiation. C) Diameter of organoids in each condition. Bars represent median and interquartile range. *p<0.05, p**<0.01, p****<0.0001 using two-way ANOVA followed by Sidak’s multiple comparisons test. N = 3 independent differentiations. D) Cell yield normalized to volume of BsM used during culture for embedding organoids or as matrix microcarriers. Bars represent mean and SEM. No statistical significance using Welch’s t-test. N = 3 independent differentiations. E) WNT9B concentration in medium normalized to cell density in each condition measured by ELISA. Bars represent mean and SEM. *p<0.05, p**<0.01 using Welch’s t-test. N = 3 independent differentiations.

**Supplemental Figure 14.**
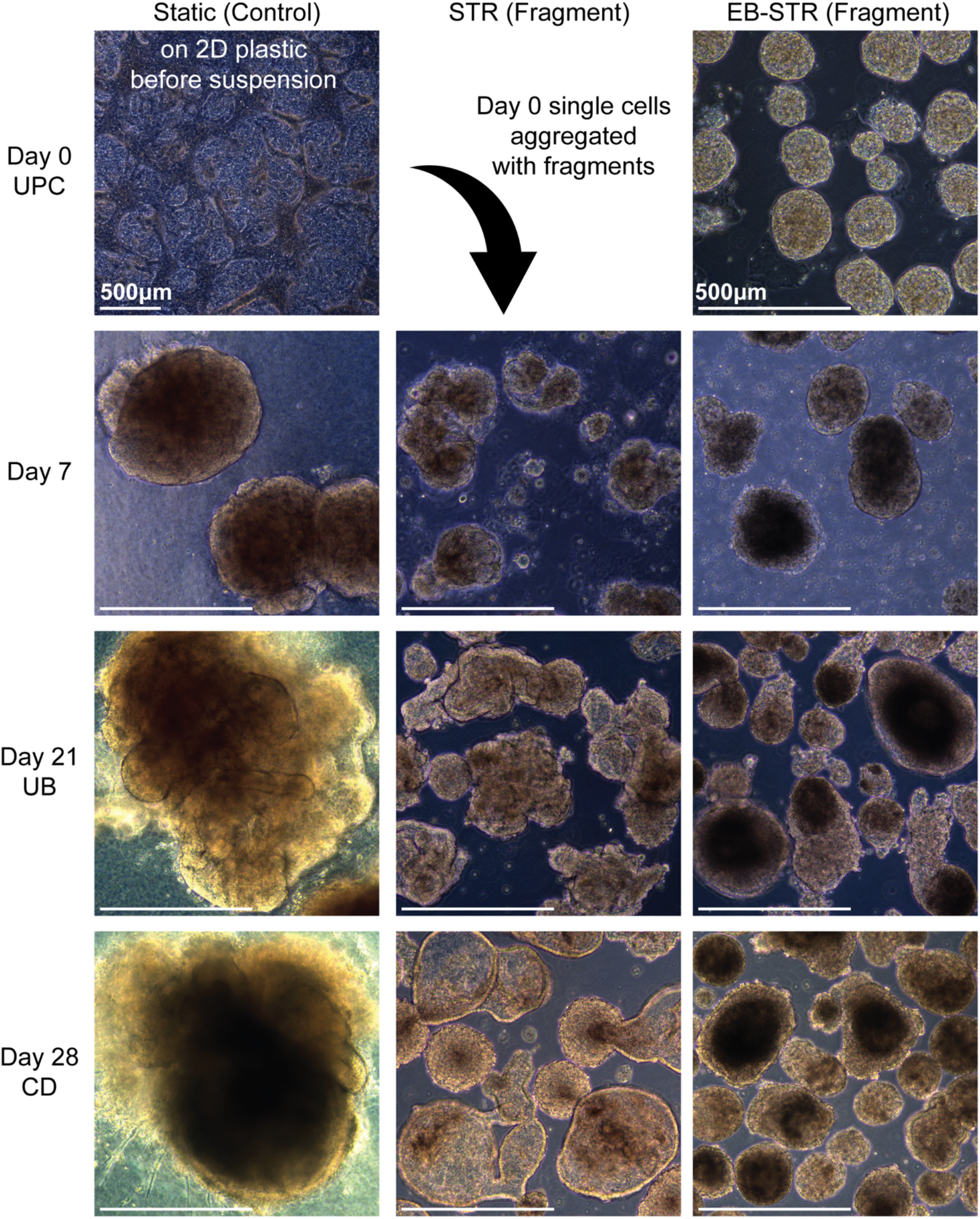
Time course of UB and CD organoids (brightfield images) on orbital shakers derived from UPCs or EBs and static embedded controls. Scale = 500 μm. N = 3 independent differentiations, all images shown from the same differentiation.

**Supplemental Figure 15.**
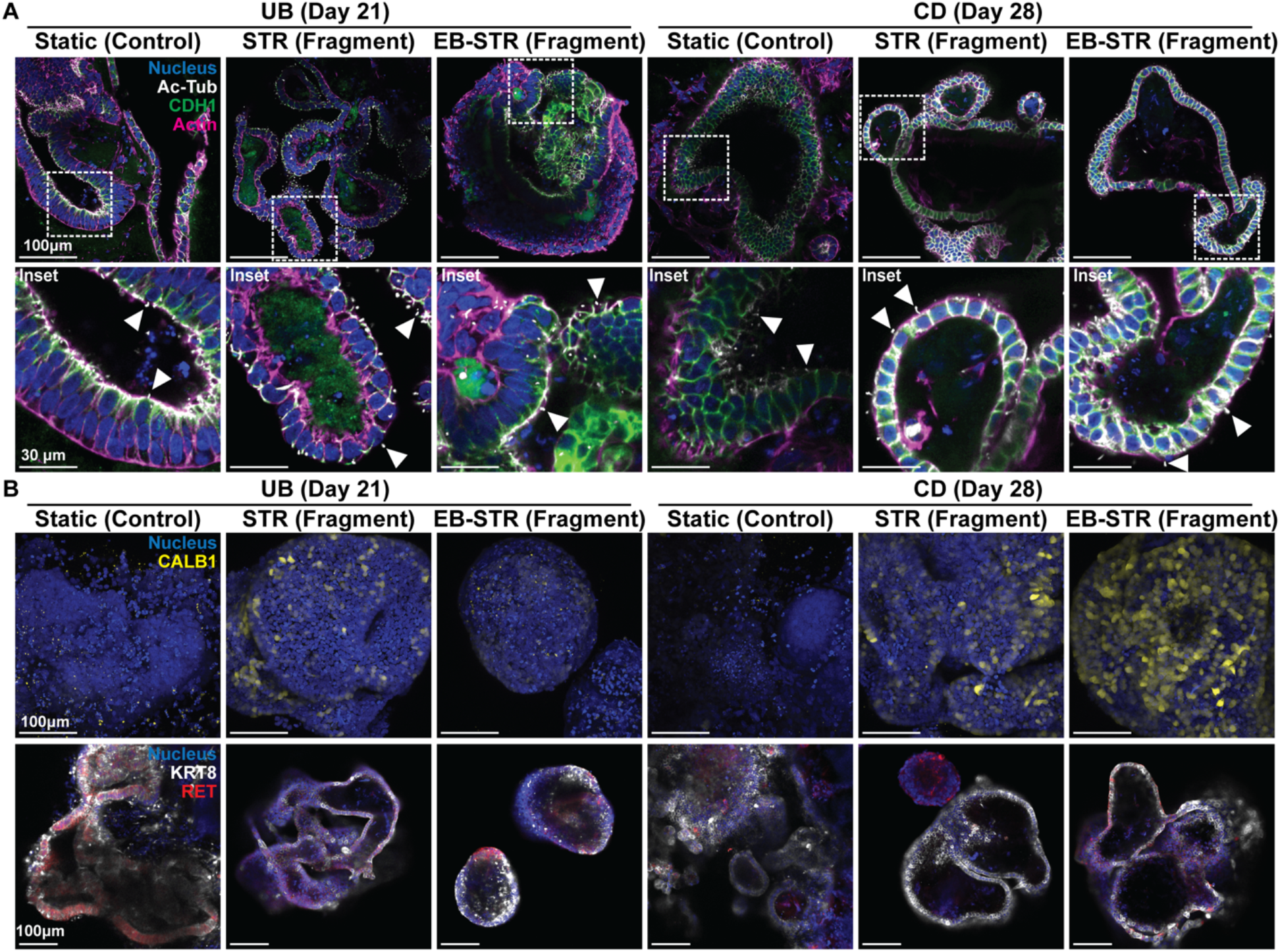
Characterization of UB and CD organoid morphology and marker expression in STR conditions and static embedded controls. Data is depicted from Figure 3, here including additional data on EB-STR (Fragment) for clarity. A) Representative immunofluorescence images of organoids in control and STR conditions. White box indicates inset. Blue = nucleus, white = Ac-Tub, green = CDH1, magenta = actin. Images are single slices. Scale = 100 μm. Inset shows apical surfaces oriented internally or externally. White arrows mark single primary cilium and apical surface. Scale = 30 μm. B) Representative immunofluorescence images of organoids in control and STR conditions. Blue = nucleus, yellow = CALB1, white = KRT8, red = RET. Images are maximum intensity projections (CALB1) or single slices (KRT8/RET). Scale = 100 μm. N = 3 independent differentiations, all images shown from the same differentiation.

**Supplemental Figure 16.**
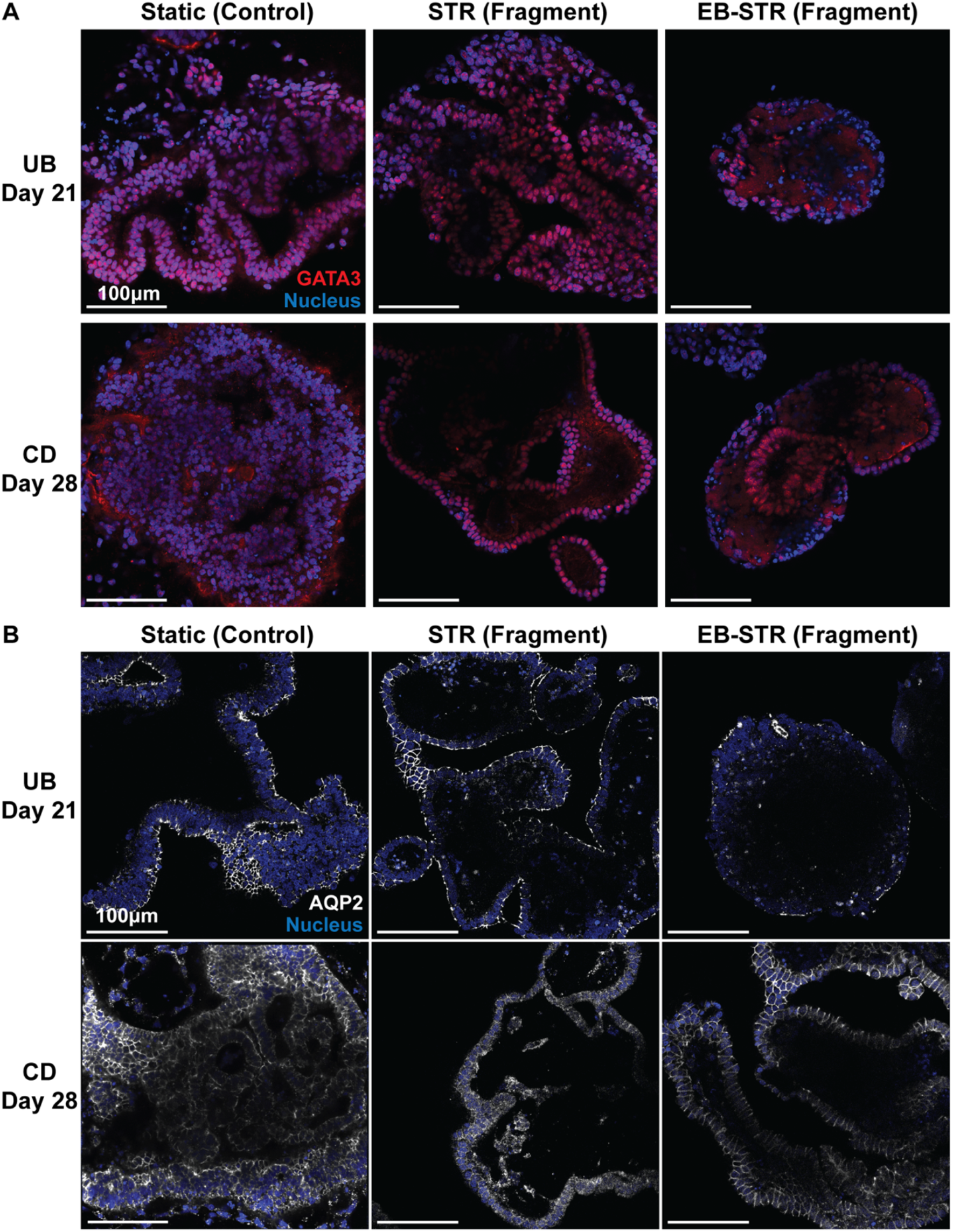
Characterization of marker expression of UB and CD organoids in STR bioreactors and cell culture inserts. A) Immunofluorescence images (single slices) of STR and static organoids at 21 days (UB) and 28 days (CD) in culture. Red = GATA3, blue = nucleus. Scale = 100 μm. B) Immunofluorescence images (single slices) of STR and static organoids at 21 days (UB) and 28 days (CD) in culture. White = AQP2, blue = nucleus. Scale = 100 μm. N = 3 independent differentiations, all images shown from the same differentiation.

**Supplemental Figure 17.**
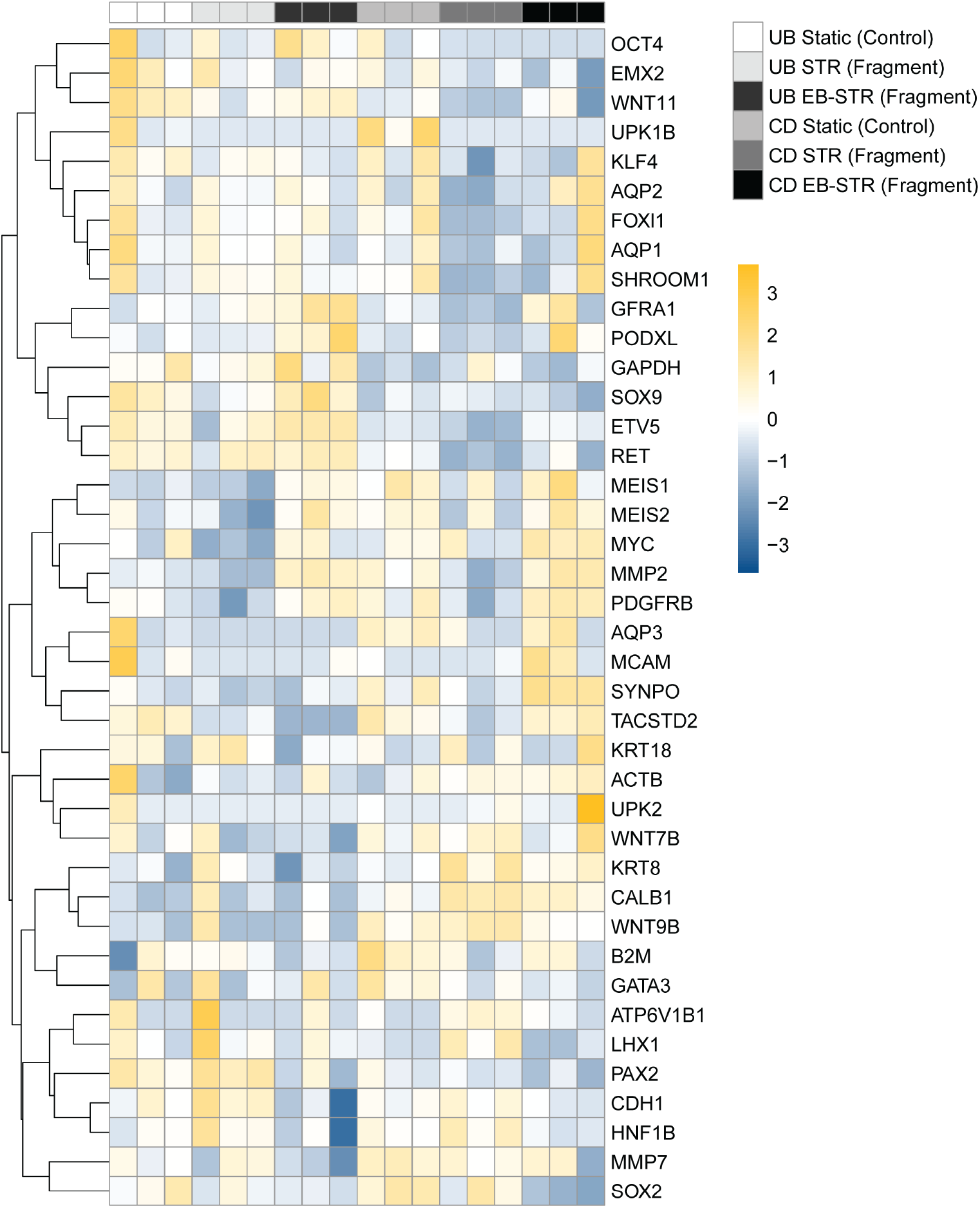
Transcriptomic analysis of UB and CD organoids derived from UPCs and EBs with matrix microcarriers on orbital shakers and static embedded controls. All tested genes above threshold are shown. N = 3 independent differentiations.

**Supplemental Figure 18.**
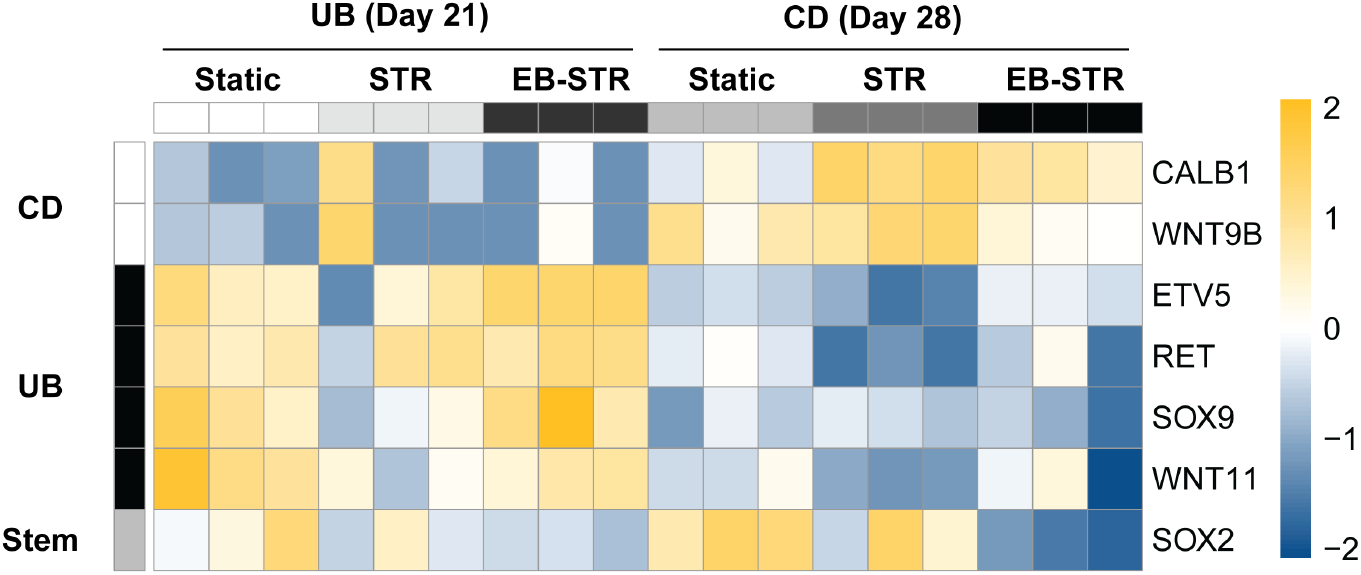
Transcriptomic analysis of UB and CD organoids derived from UPCs and EBs with matrix microcarriers on orbital shakers and static embedded controls. Key UB and CD markers are shown. N = 3 independent differentiations.

**Supplemental Movie 1.**
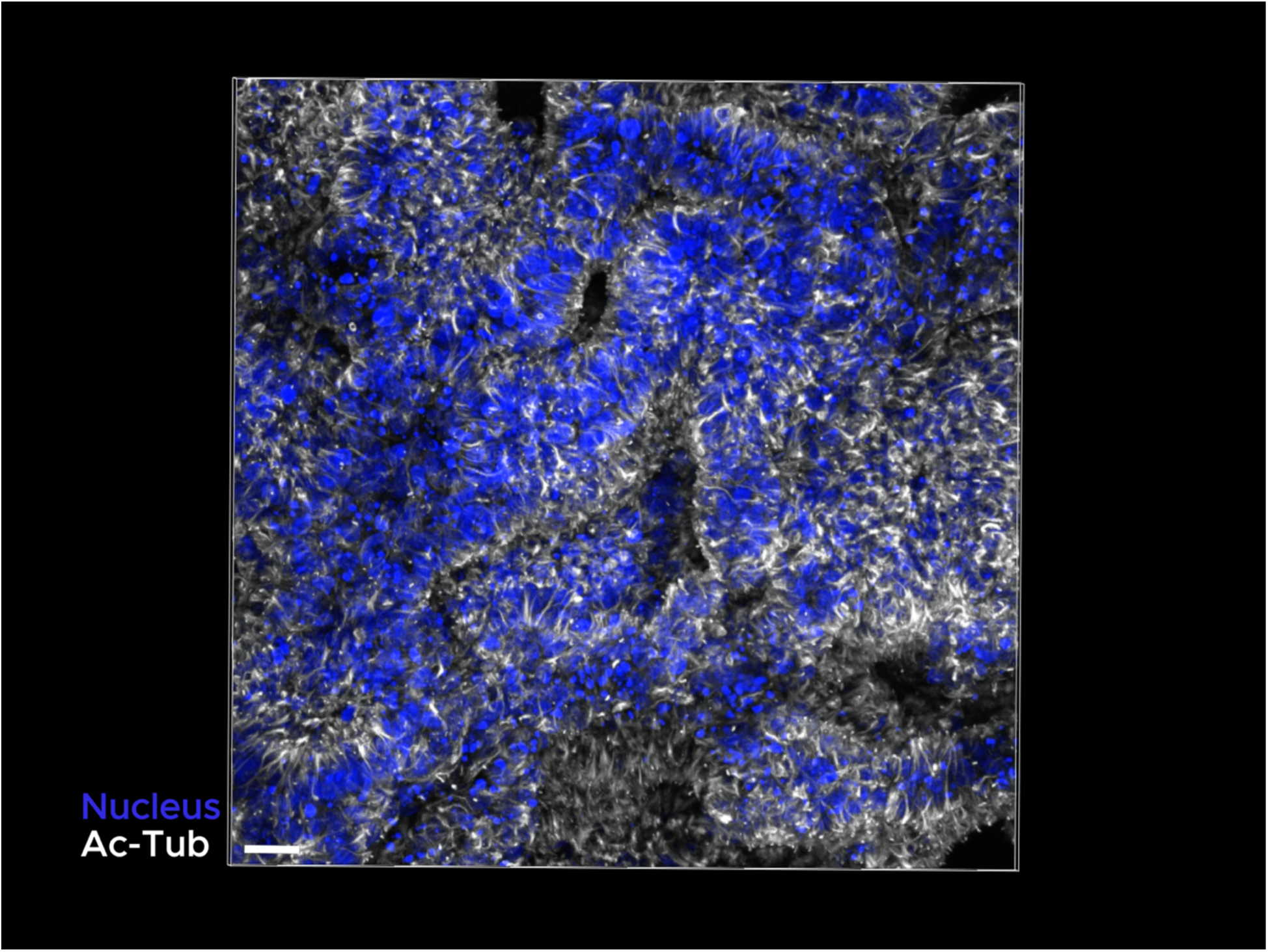
Matrix microcarriers for UB organoid culture on orbital shakers. Perspective view of UB organoid surface cultured on fragment microcarriers on an orbital shaker after 21 days in culture, showing apically-oriented cilia in lumens that are contiguous with surface. Blue = nucleus, white = Ac-Tub. Scale = 30 μm.

**Table S1.**
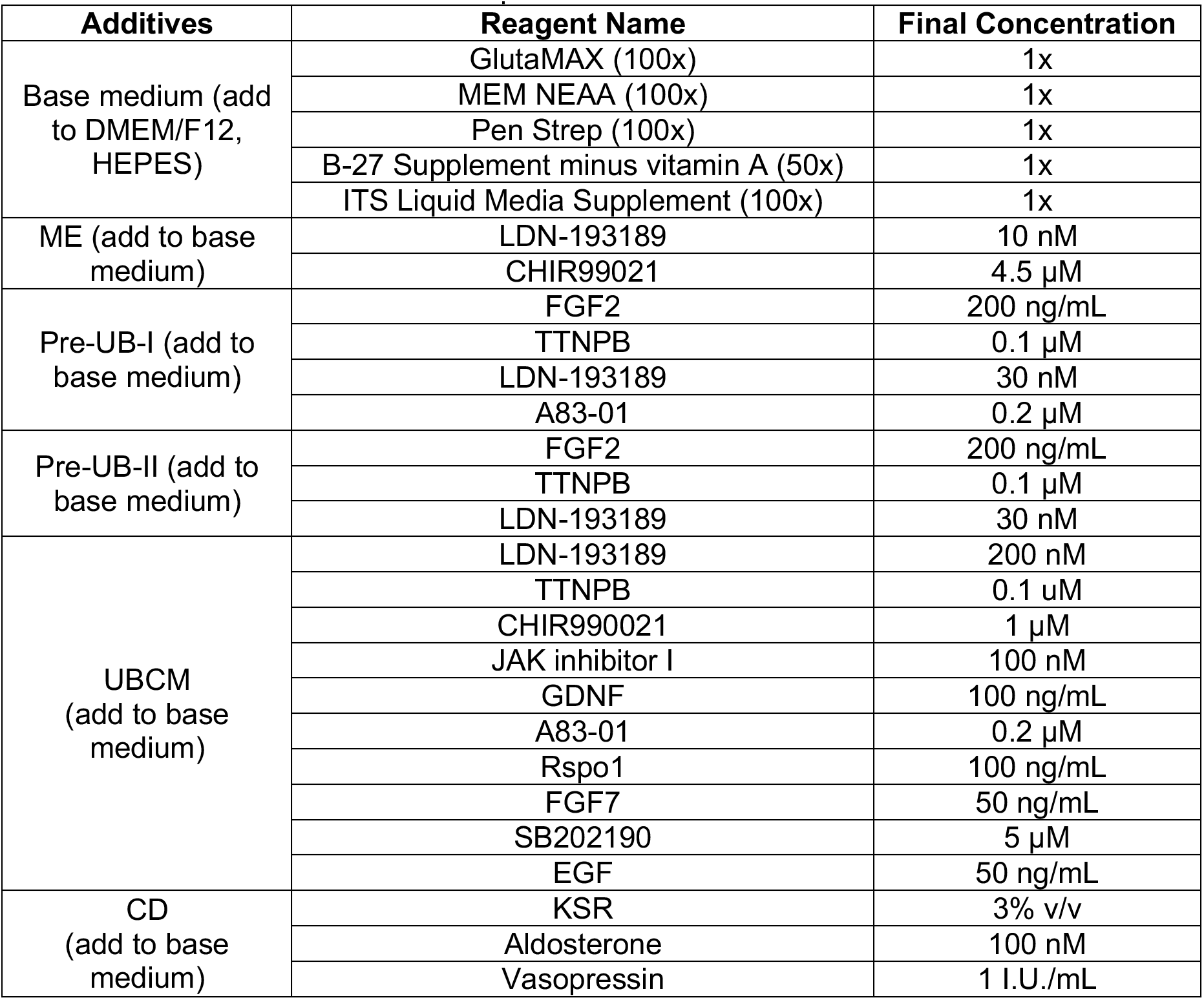
Differentiation medium compositions.

**Table S2.**
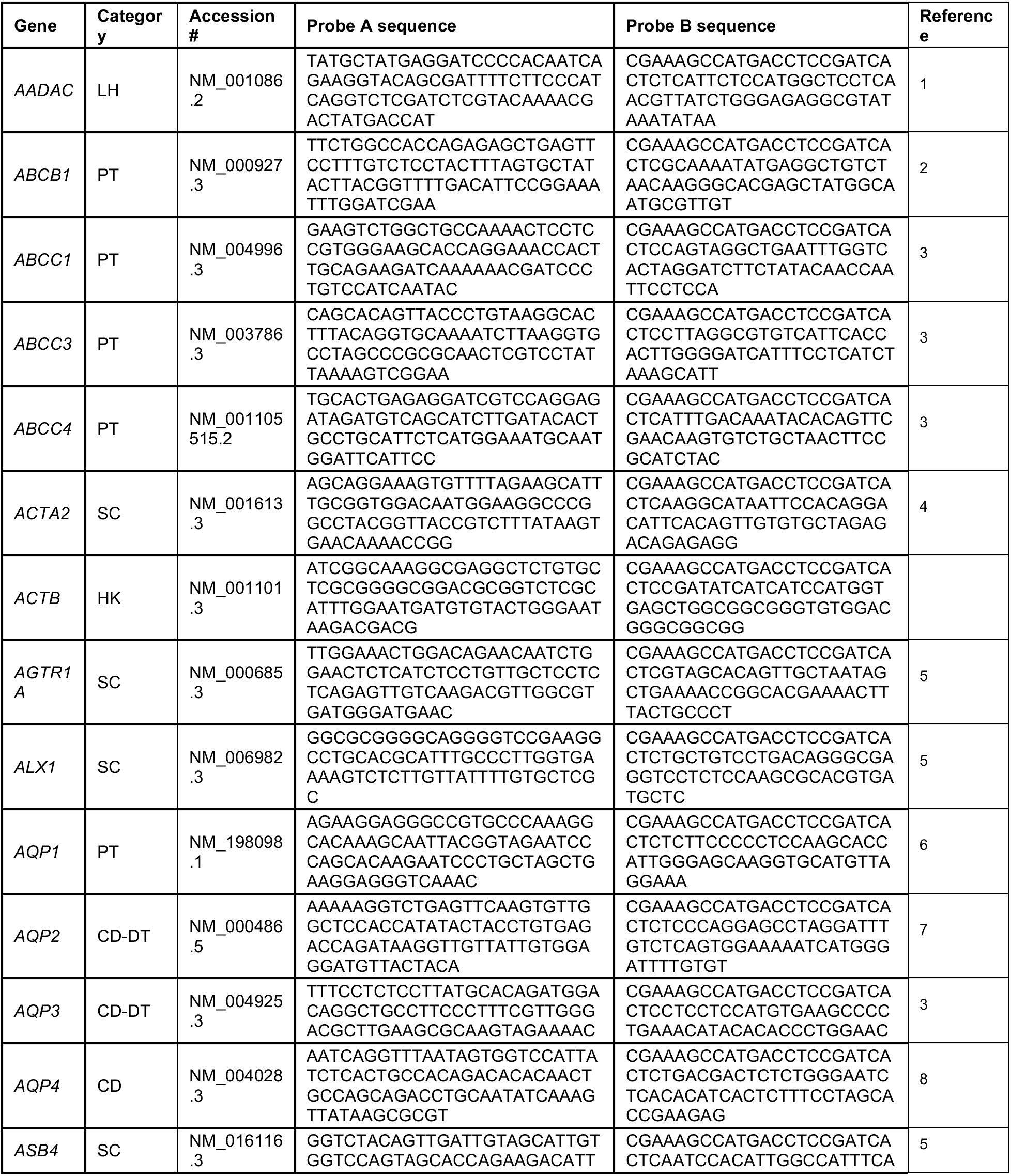
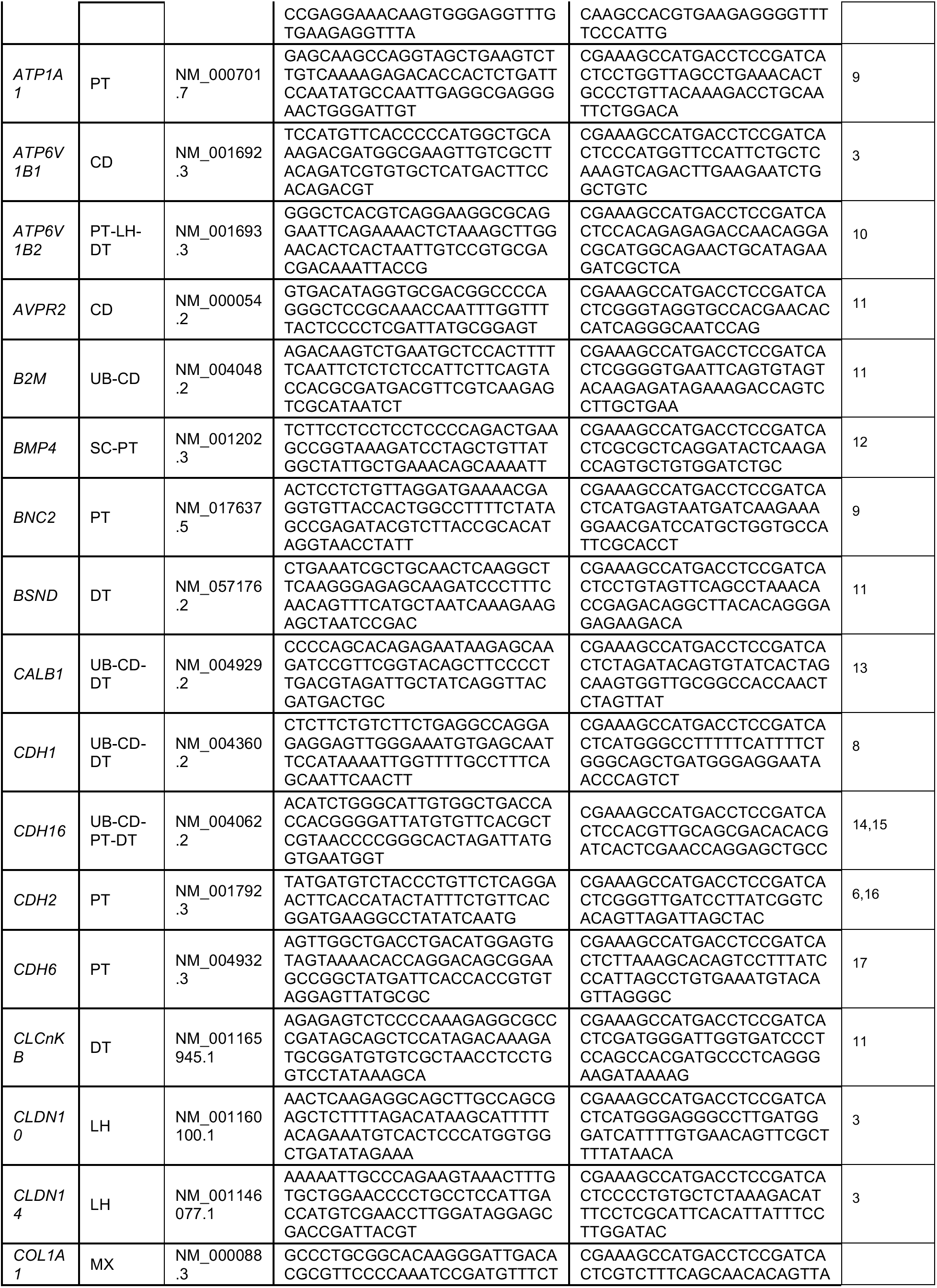
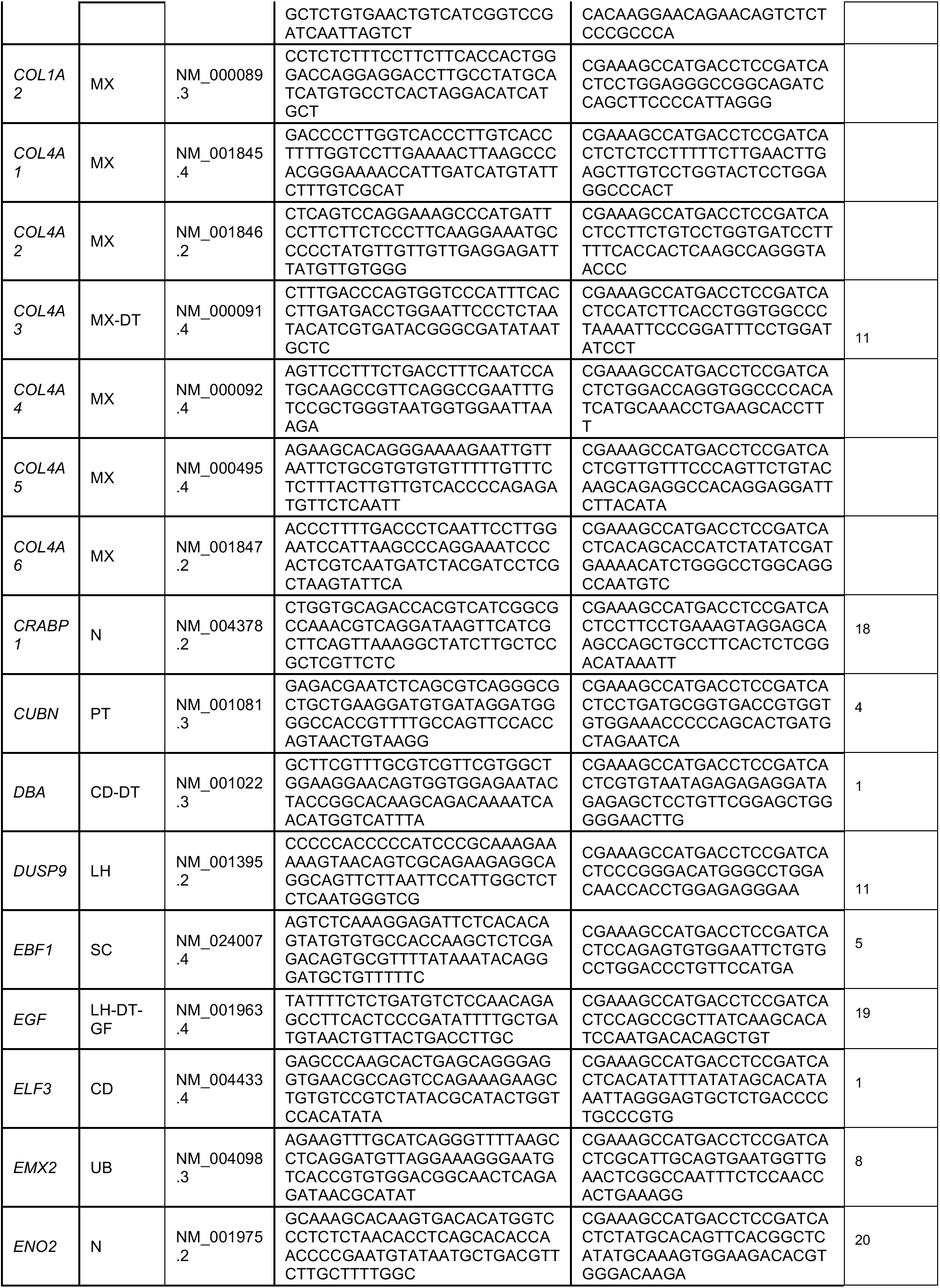
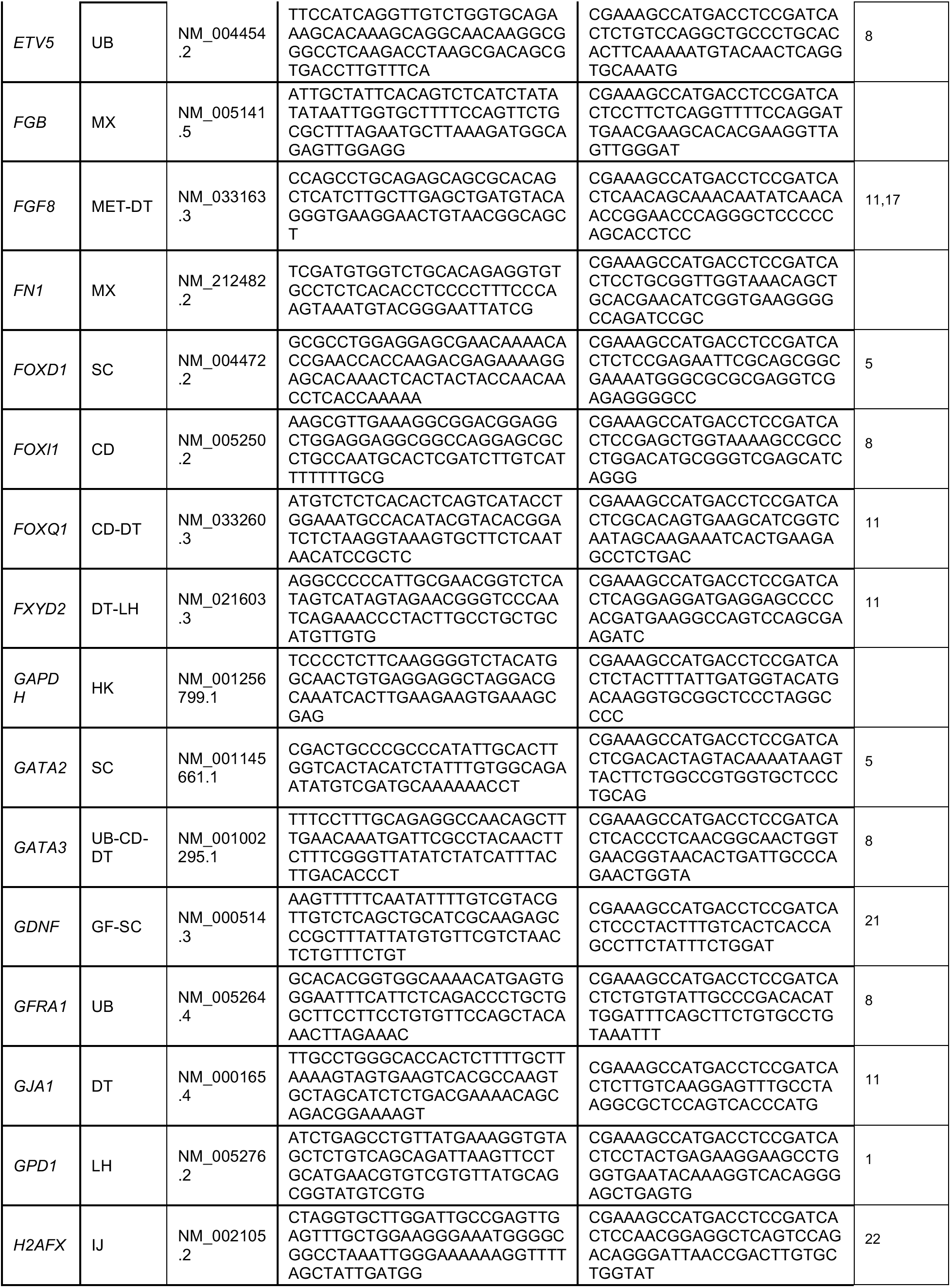
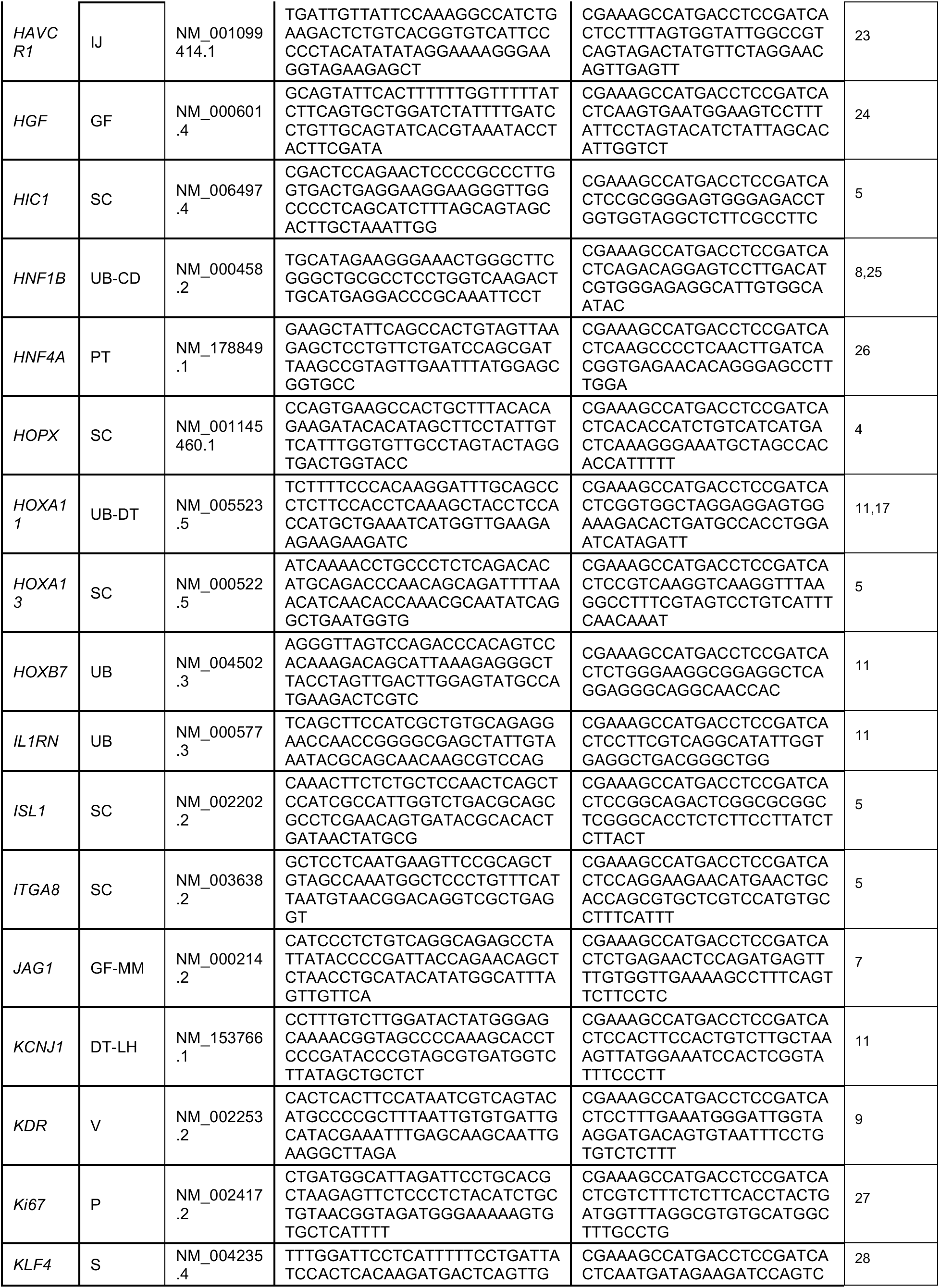
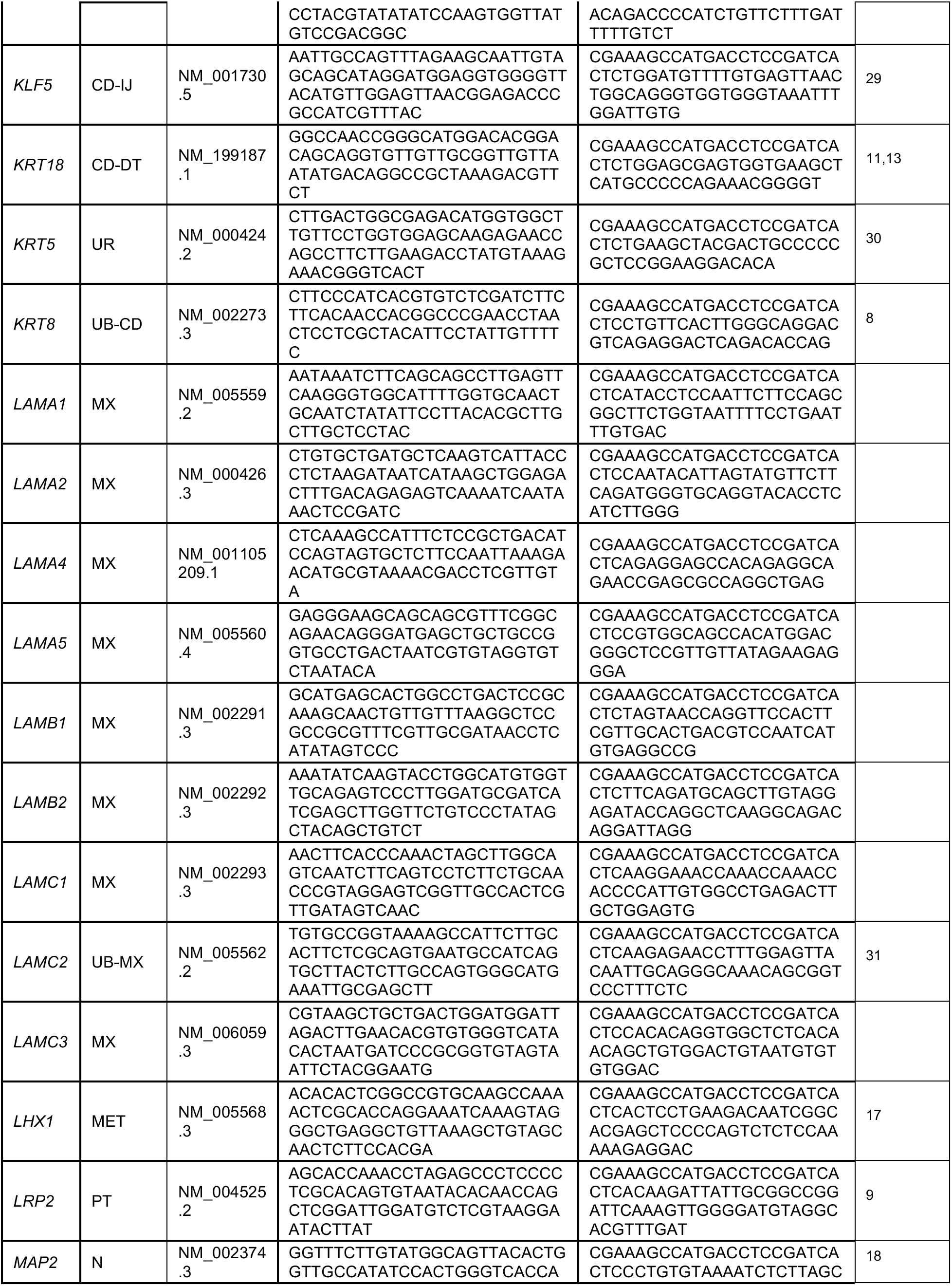
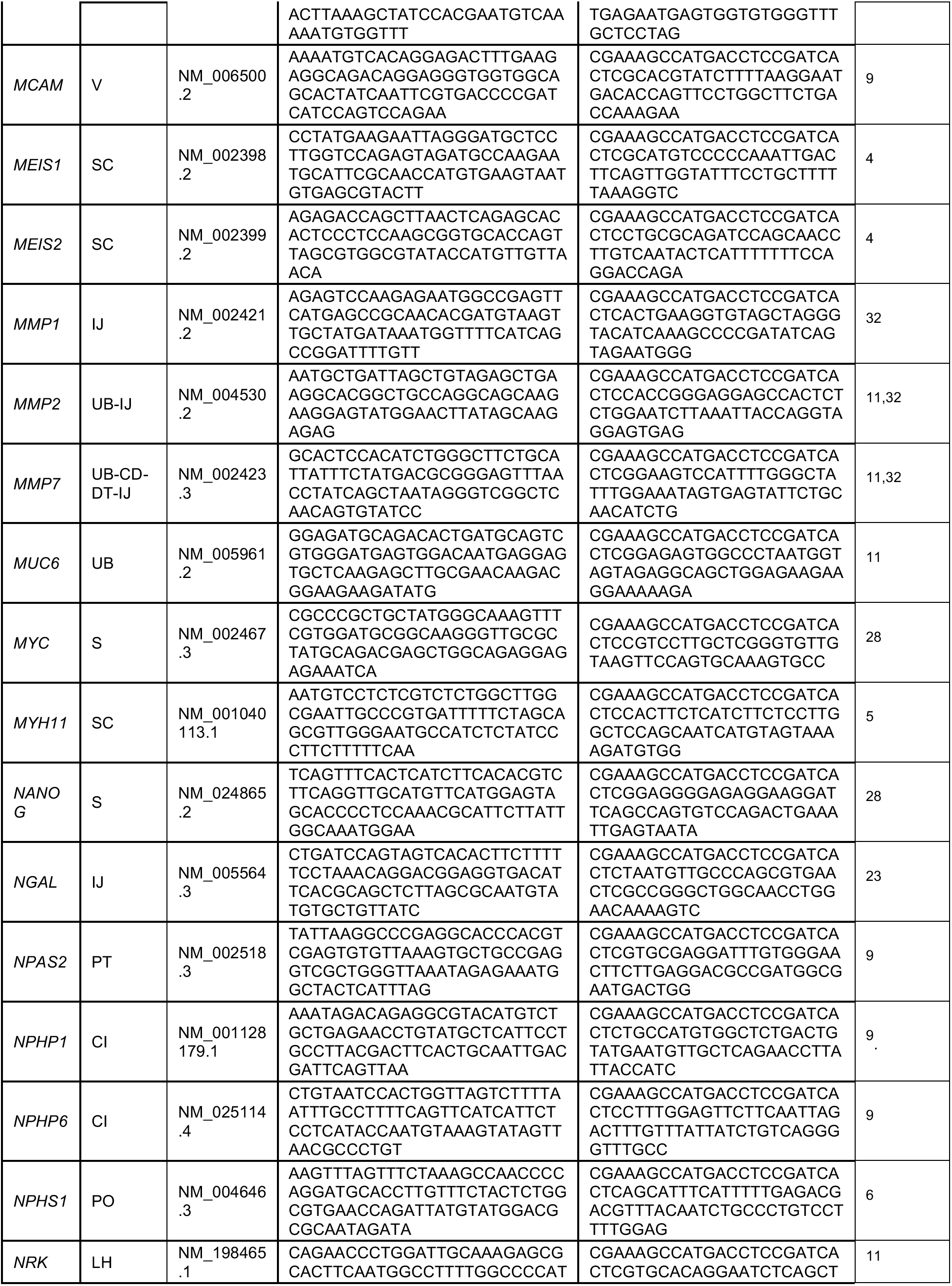
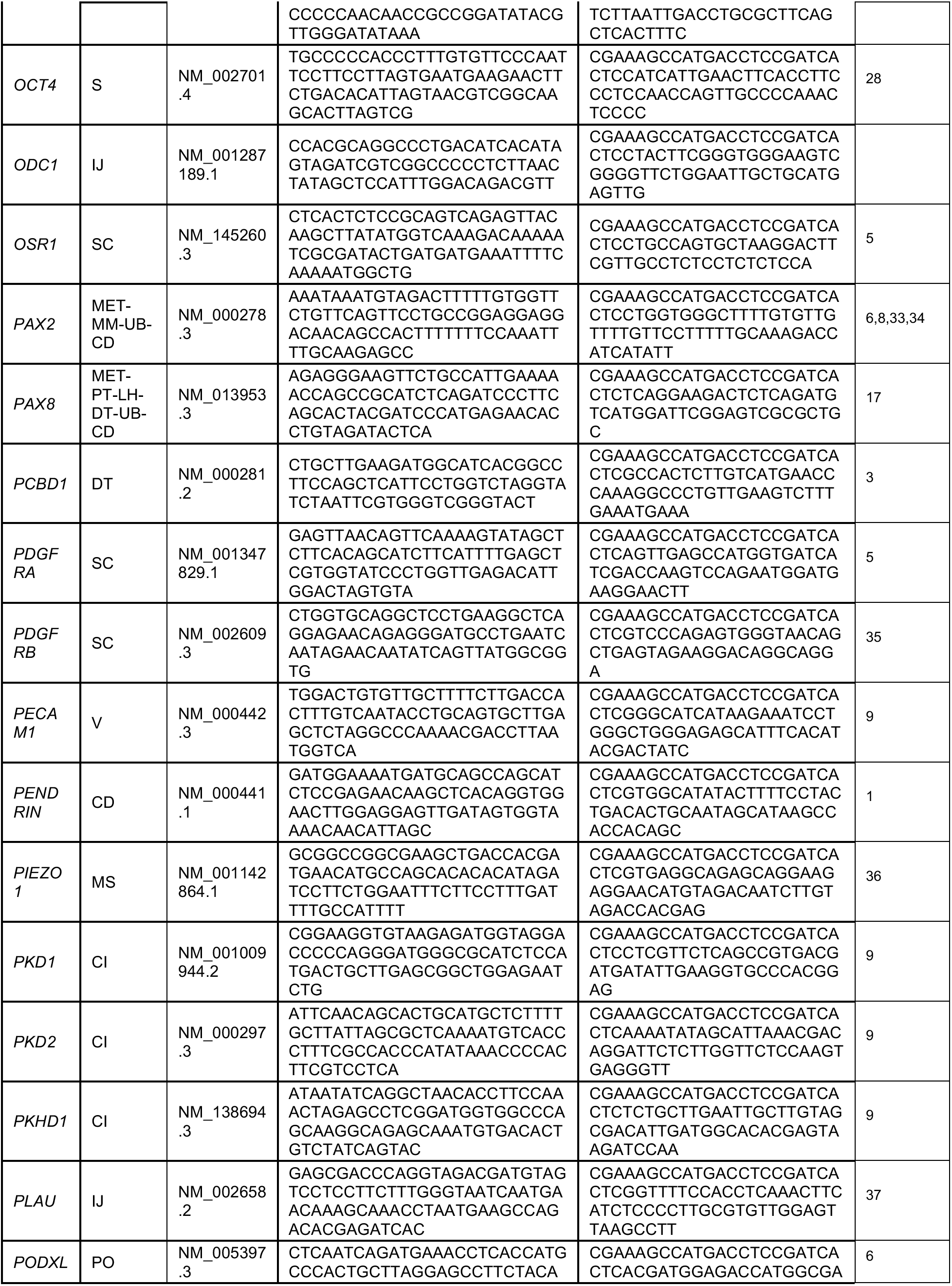
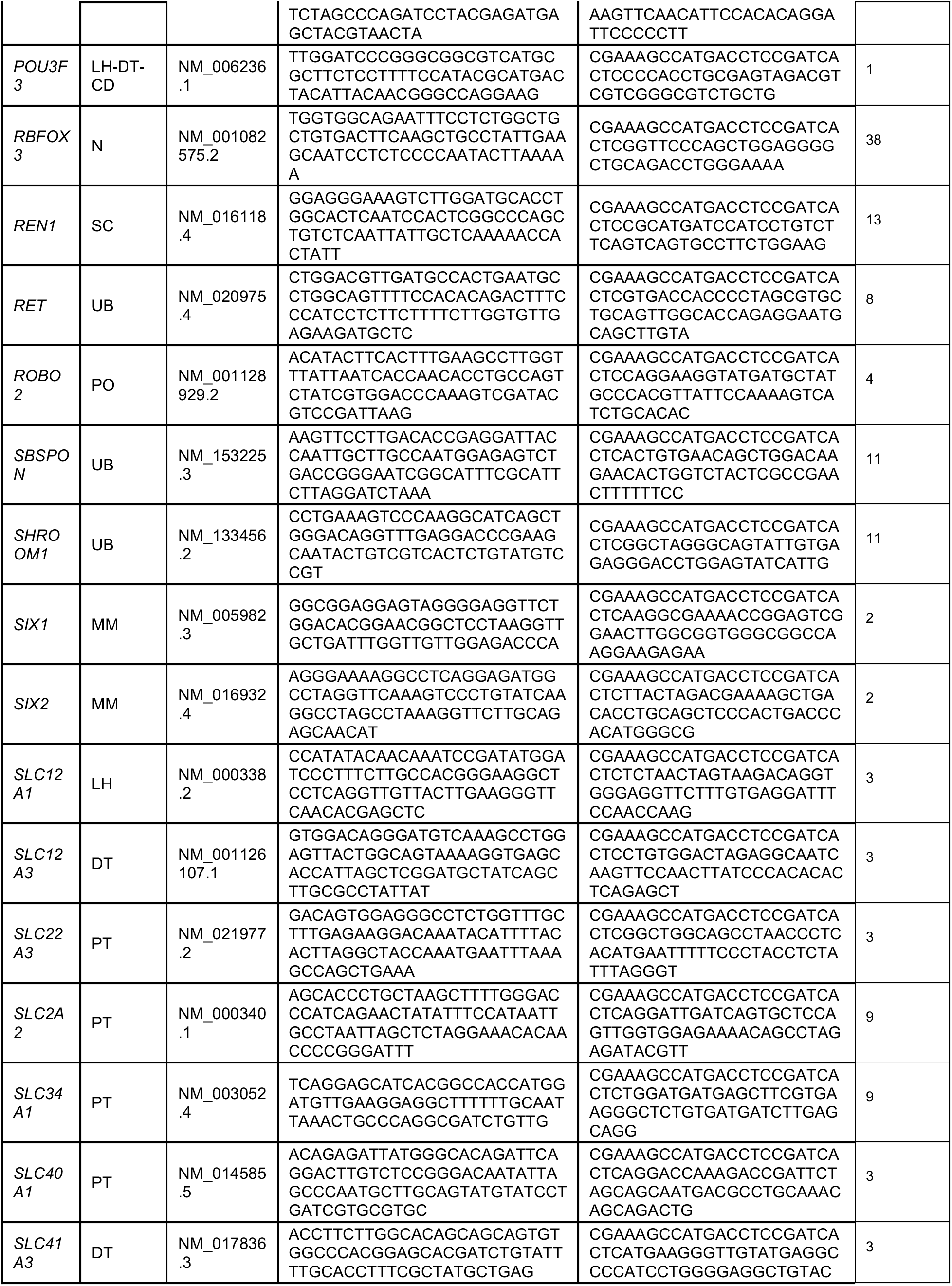
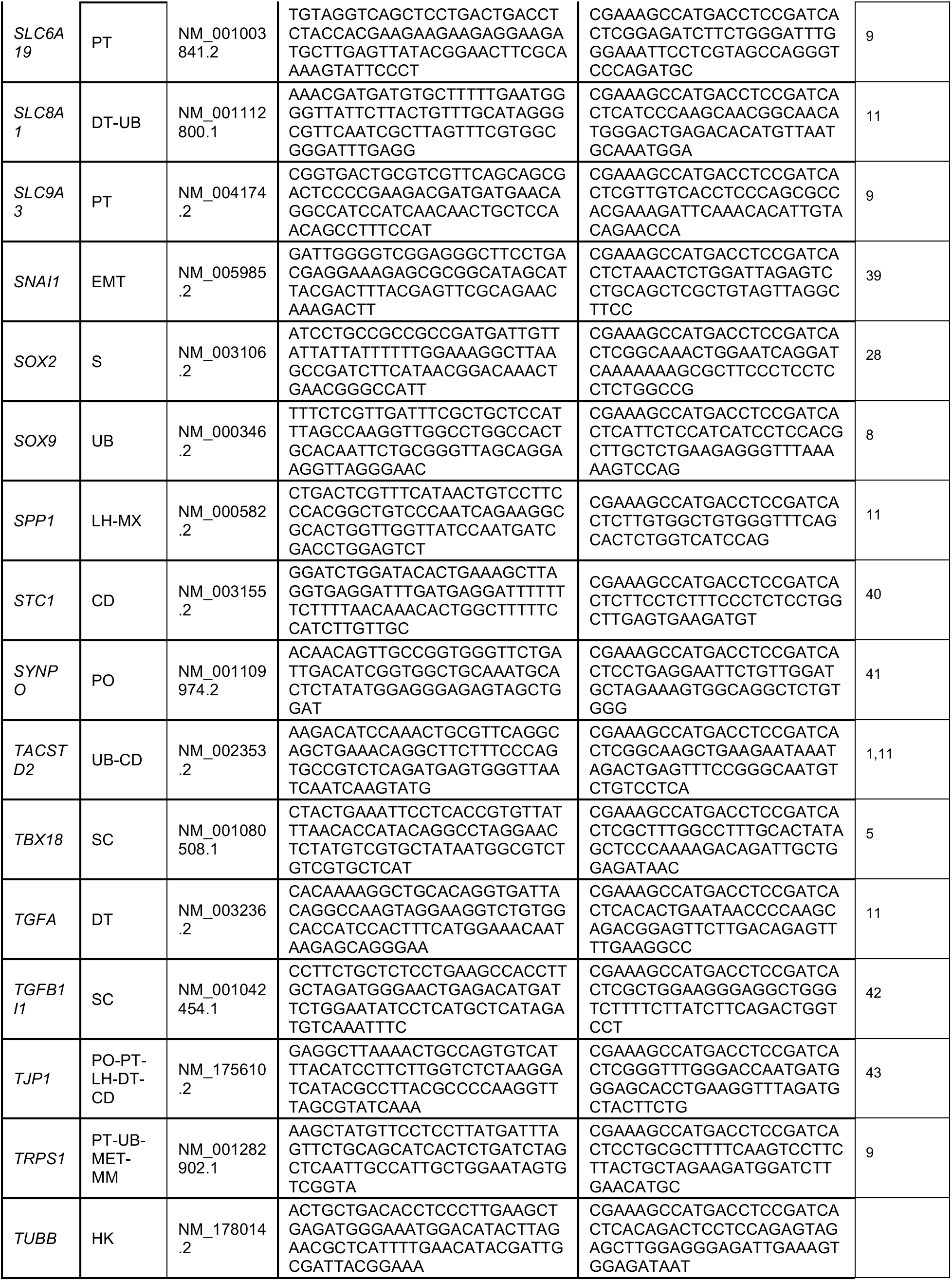
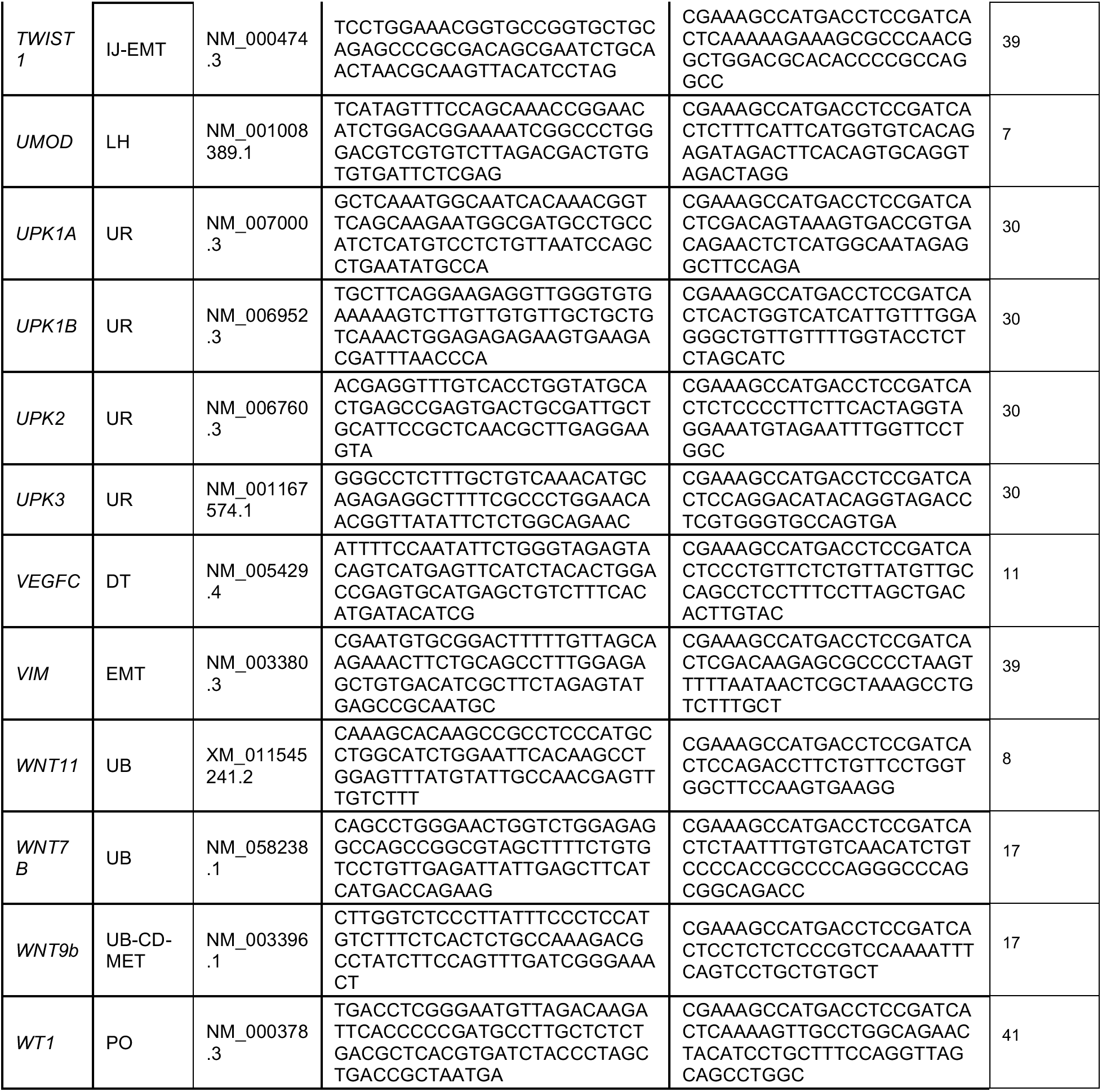
Gene panel for Nanostring transcriptomic analysis. Table includes gene name, accession number, marker or functional categorization, probe sequences, and references used for categorization. CD = collecting duct, Cl = Cilia, DT = distal tubule, EMT = epithelial to mesenchymal transition, GF = growth factor, HK = Housekeeping, IJ = injury, LH = loop of henle, MET = mesenchymal to epithelial transition, MM = metanephric mesenchyme, MS = mechanosensitive, MX = matrix, N = neural, P = proliferation, PO = podocyte, PT = proximal tubule, S = stem cell, SC = stromal cell, UR = ureter, V = vasculature.

**Table S3.**
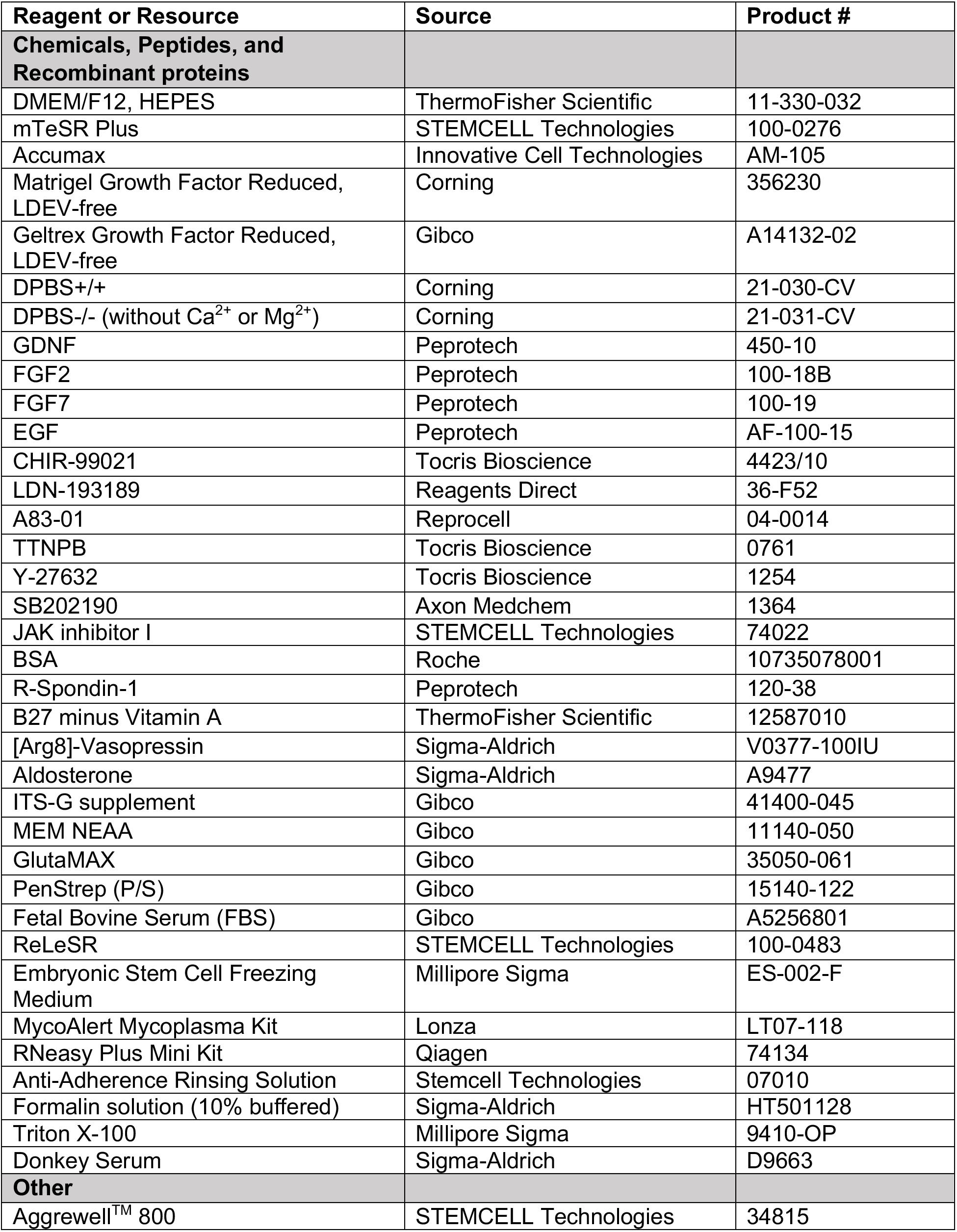
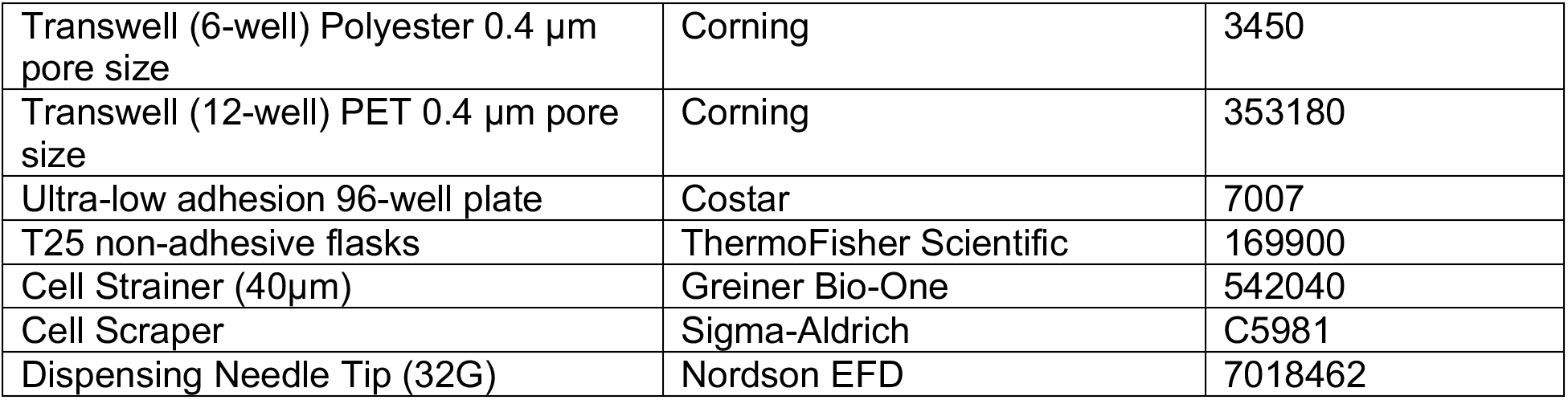
Resources and Reagents.

**Table S4.** Antibodies and Stains.

| Target | Host Species | Manufacturer | Product # | Dilution |
| --- | --- | --- | --- | --- |
| **Primaries** |  |  |  |  |
| KRT8 | Rat | DSHB | TROMA-1 | 1:50 |
| E-Cadherin | Mouse | Abcam | ab1416 | 1:100 |
| ETV5 | Rabbit | Abcam | ab102010 | 1:400 |
| SOX9 | Rabbit | Abcam | ab185230 | 1:300 |
| PAX2 | Rabbit | ThermoFisher Scientific | 71-6000 | 1:200 |
| GATA3 | Rabbit | Cell Signaling | 5852T | 1:100 |
| ATP6V1B1 | Rabbit | Abcam | ab192612 | 1:200 |
| AQP2 | Rabbit | Abcam | ab62628 | 1:200 |
| ENaC | Rabbit | Invitrogen | PA1-920A | 1:200 |
| Na+K+ATPase | Rabbit | Abcam | ab76020 | 1:200 |
| RET | Goat | R&D technologies | AF1486 | 1:100 |
| CD117 | Mouse | BioLegend | 313204 | 1:300 |
| Acetylated Tubulin | Rabbit | Cell Signaling Technologies | 5335 | 1:200 |
| CALB1 | Rabbit | Abcam | ab108404 | 1:200 |
| **Secondaries** |  |  |  |  |
| 488 anti Rat | Donkey | ThermoFisher Scientific | A21208 | 1:400 |
| 546 anti Mouse | Donkey | ThermoFisher Scientific | A10036 | 1:400 |
| 647 anti Rabbit | Donkey | ThermoFisher Scientific | A32795 | 1:300 |
| 647 anti Rat | Donkey | Abcam | ab150155 | 1:200 |
| 555+ anti Rabbit | Donkey | ThermoFisher Scientific | A32794 | 1:300 |
| 488 anti Mouse | Donkey | ThermoFisher Scientific | A21202 | 1:300 |
| 546 anti Goat | Donkey | ThermoFisher Scientific | A11056 | 1:300 |
| **Cell stains** |  |  |  |  |
| DAPI (4',6-diamidino-2-phenylindole) |  | ThermoFisher Scientific | D1306 | 1:1000 |
| ActinGreen 488 ReadyProbes |  | ThermoFisher Scientific | R37110 | 1:100 |

## References

1. Yousef Yengej, F. A., Jansen, J., Rookmaaker, M. B., Verhaar, M. C. & Clevers, H. Kidney Organoids and Tubuloids. Cells 9, 1–20 (2020).

2. McCormick, F., Held, P. J. & Chertow, G. M. The terrible toll of the kidney shortage. Journal of the American Society of Nephrology 29, 2775–2776 (2018).

3. Woolf, A. S. Growing a new human kidney. Kidney International vol. 96 871–882 (2019).

4. Ahmed, S. M., Shivnaraine, R. V. & Wu, J. C. FDA Modernization Act 2.0 Paves the Way to Computational Biology and Clinical Trials in a Dish. Circulation 148, 309–311 (2023).

5. Wolf, K. J., Weiss, J. D., Uzel, S. G. M., Skylar-Scott, M. A. & Lewis, J. A. Biomanufacturing human tissues via organ building blocks. Cell Stem Cell 29, 667–677 (2022).

6. McMahon, A. P. Development of the Mammalian Kidney. Curr. Top. Dev. Biol. 117, 31–64 (2016).

7. Huang, B. et al. Spatially patterned kidney assembloids recapitulate progenitor self-assembly and enable high-fidelity in vivo disease modeling. Cell Stem Cell (2025).

8. Tanigawa, S. et al. Generation of the organotypic kidney structure by integrating pluripotent stem cell-derived renal stroma. Nat. Commun. 13, 1–15 (2022).

9. Shi, M. et al. Integrating collecting systems in human kidney organoids through fusion of distal nephron to ureteric bud. Cell Stem Cell 32, 1–16 (2025).

10. Lawlor, K. T. et al. Cellular extrusion bioprinting improves kidney organoid reproducibility and conformation. Nat. Mater. 20, 260–271 (2020).

11. Skylar-Scott, M. A. et al. Biomanufacturing of organ-specific tissues with high cellular density and embedded vascular channels. Sci. Adv. 5, eaaw2459 (2019).

12. Brassard, J. A., Nikolaev, M., Hübscher, T., Hofer, M. & Lutolf, M. P. Recapitulating macro-scale tissue self-organization through organoid bioprinting. Nat. Mater. 20, 22–29 (2021).

13. Hatton, I. A. et al. The human cell count and size distribution. Proceedings of the National Academy of Sciences 120, e2303077120 (2023).

14. Bianconi, E. et al. An estimation of the number of cells in the human body. Ann. Hum. Biol. 40, 463–471 (2013).

15. Hirokawa, Y. et al. Low-viscosity matrix suspension culture enables scalable analysis of patient-derived organoids and tumoroids from the large intestine. *Commun*. Biol. 4, 1–17 (2021).

16. Takahashi, J. et al. Suspension culture in a rotating bioreactor for efficient generation of human intestinal organoids. Cell Reports Methods 2, 100337 (2022).

17. Schneeberger, K. et al. Large-Scale Production of LGR5-Positive Bipotential Human Liver Stem Cells. Hepatology 72, 257–270 (2020).

18. Co, J. Y., Klein, J. A., Kang, S. & Homan, K. A. Suspended hydrogel culture as a method to scale up intestinal organoids. Sci. Rep. 13, 1–12 (2023).

19. Morizane, R. & Bonventre, J. V. Generation of nephron progenitor cells and kidney organoids from human pluripotent stem cells. Nat. Protoc. 12, 195–207 (2017).

20. Takasato, M., Er, P. X., Chiu, H. S. & Little, M. H. Generation of kidney organoids from human pluripotent stem cells. Nat. Protoc. 11, 1681–1692 (2016).

21. Zeng, Z. & Huang, B. et al. Generation of patterned kidney organoids that recapitulate the adult kidney collecting duct system from expandable ureteric bud progenitors. Nat. Commun. 12, 3614 (2021).

22. Shi, M., Fu, P., Bonventre, J. V. & McCracken, K. W. Directed differentiation of ureteric bud and collecting duct organoids from human pluripotent stem cells. Nat. Protoc. 18, 2485–2508 (2023).

23. Mae, S.-I., Ryosaka, M. & Osafune, K. Protocol to Generate Ureteric Bud Structures from Human iPS Cells. in Methods in Molecular Biology 117–123 (Humana Press, 2019).

24. Uchimura, K., Wu, H., Yoshimura, Y. & Humphreys, B. D. Human Pluripotent Stem Cell-Derived Kidney Organoids with Improved Collecting Duct Maturation and Injury Modeling. Cell Rep. 33, 108514 (2020).

25. Kumar, S. V. et al. Kidney micro-organoids in suspension culture as a scalable source of human pluripotent stem cell-derived kidney cells. Development (Cambridge) 146, (2019).

26. Sander, V. et al. Protocol for Large-Scale Production of Kidney Organoids from Human Pluripotent Stem Cells. STAR Protoc. 1, 100150 (2020).

27. Przepiorski, A. et al. A Simple Bioreactor-Based Method to Generate Kidney Organoids from Pluripotent Stem Cells. Stem Cell Reports 11, 470–484 (2018).

28. Shi, M. et al. Human ureteric bud organoids recapitulate branching morphogenesis and differentiate into functional collecting duct cell types. Nat. Biotechnol. 41, 252–261 (2022).

29. Mae, S. I. et al. Expansion of Human iPSC-Derived Ureteric Bud Organoids with Repeated Branching Potential. Cell Rep. 32, 107963 (2020).

30. Ho, D. L. L. et al. Large-Scale Production of Wholly Cellular Bioinks via the Optimization of Human Induced Pluripotent Stem Cell Aggregate Culture in Automated Bioreactors. Adv. Healthc. Mater. 11, (2022).

31. Nauli, S. M. et al. Polycystins 1 and 2 mediate mechanosensation in the primary cilium of kidney cells. Nat. Genet. 33, 129–137 (2003).

32. Resnick, A. & Hopfer, U. Force-response considerations in ciliary mechanosensation. Biophys. J. 93, 1380–1390 (2007).

33. Weinbaum, S., Duan, Y., Satlin, L. M., Wang, T. & Weinstein, A. M. Mechanotransduction in the renal tubule. Am. J. Physiol. Renal Physiol. 299, 1220–1236 (2010).

34. Weinbaum, S., Duan, Y., Thi, M. M. & You, L. An integrative review of mechanotransduction in endothelial, epithelial (renal) and dendritic cells (osteocytes). Cell. Mol. Bioeng. 4, 510–537 (2011).

35. Flitney, E. W., Kuczmarski, E. R., Adam, S. A. & Goldman, R. D. Insights into the mechanical properties of epithelial cells: the effects of shear stress on the assembly and remodeling of keratin intermediate filaments. The FASEB Journal 23, 2110–2119 (2009).

36. Molladavoodi, S., Robichaud, M., Wulff, D. & Gorbet, M. Corneal epithelial cells exposed to shear stress show altered cytoskeleton and migratory behaviour. PLoS One 12, 1–16 (2017).

37. Wu, H. et al. Comparative Analysis and Refinement of Human PSC-Derived Kidney Organoid Differentiation with Single-Cell Transcriptomics. Cell Stem Cell 23, 869–881.e8 (2018).

38. Kelm, J. M., Timmins, N. E., Brown, C. J., Fussenegger, M. & Nielsen, L. K. Method for generation of homogeneous multicellular tumor spheroids applicable to a wide variety of cell types. Biotechnol. Bioeng. 83, 173–180 (2003).

39. Liu, D., Chen, S. & Naing, M. W. A review of manufacturing capabilities of cell spheroid generation technologies and future development. Biotechnol. Bioeng. 118, 542–554 (2021).

40. Wolf, K. J. et al. Perfusable 3D models of ureteric bud and collecting duct tubules. Cell Biomaterials 2, 100297 (2026).

41. Carroll, T. J., Park, J., Hayashi, S., Majumdar, A. & Mcmahon, A. P. Wnt9b Plays a Central Role in the Regulation of Mesenchymal to Epithelial Transitions Underlying Organogenesis of the Mammalian Urogenital System. 9, 283–292 (2005).

42. Liu, W. et al. Effect of flow and stretch on the [Ca2+]i response of principal and intercalated cells in cortical collecting duct. Am. J. Physiol. Renal Physiol. 285, 998–1012 (2003).

43. Liu, W., Morimoto, T., Woda, C., Kleyman, T. R. & Satlin, L. M. Ca2+ dependence of flow-stimulated K secretion in the mammalian cortical collecting duct. Am. J. Physiol. Renal Physiol. 293, 227–235 (2007).

44. Woda, C. B. et al. Ontogeny of flow-stimulated potassium secretion in rabbit cortical collecting duct: Functional and molecular aspects. Am. J. Physiol. Renal Physiol. 285, 629–639 (2003).

45. Evan, A., Satlin, L., Gattone, V., Connors, B. & Schwartz, G. Postnatal maturation of rabbit renal collecting duct. II. Morphological observations. Am. J. Physiol. Renal Physiol. 261, F91–F107 (1991).

46. Conrad, L. et al. The biomechanical basis of biased epithelial tube elongation in lung and kidney development. Development 148, 1–15 (2021).

47. Muir, V. G. et al. Influence of Microgel and Interstitial Matrix Compositions on Granular Hydrogel Composite Properties. Advanced Science 2206117, 1–17 (2023).

## Supplemental References

1. Tsujimoto, H. et al. Human Kidney Lineages from Pluripotent Stem Cells A Modular Differentiation System Maps Multiple Human Kidney Lineages from Pluripotent Stem Cells. Cell Reports 31, 107476 (2020).

2. Takasato, M. et al. Kidney organoids from human iPS cells contain multiple lineages and model human nephrogenesis. Nature 526, 564–568 (2015).

3. Schutgens, F. et al. Tubuloids derived from human adult kidney and urine for personalized disease modeling. Nat Biotechnol 37, 303–313 (2019).

4. Combes, A. N., Zappia, L., Er, P. X., Oshlack, A. & Little, M. H. Single-cell analysis reveals congruence between kidney organoids and human fetal kidney. Genome Med 11, 1–15 (2019).

5. Tanigawa, S. et al. Generation of the organotypic kidney structure by integrating pluripotent stem cell-derived renal stroma. Nat Commun 13, 1–15 (2022).

6. Morizane, R. et al. Nephron organoids derived from human pluripotent stem cells model kidney development and injury. Nat Biotechnol 33, 1193–1200 (2015).

7. Lindström, N. O. et al. Conserved and divergent features of human and mouse kidney organogenesis. Journal of the American Society of Nephrology 29, 785–805 (2018).

8. Zeng, Z. et al. Generation of patterned kidney organoids that recapitulate the adult kidney collecting duct system from expandable ureteric bud progenitors. Nat Commun 12, 3614 (2021).

9. Homan, K. A. et al. Flow-enhanced vascularization and maturation of kidney organoids in vitro. Nat Methods 16, 255–262 (2019).

10. Paunescu, T. G., Da Silva, N., Marshansky, V., McKee, M. & Brown, D. Expression of the 56-kDa B2 subunit isoform of the vacuolar H+-ATPase in proton-secreting cells of the kidney and epididymis. American Journal of Physiology-Cell Physiology 287, C149–C162 (2004).

11. Howden, S. E. et al. Plasticity of distal nephron epithelia from human kidney organoids enables the induction of ureteric tip and stalk. Cell Stem Cell 28, 1–14 (2021).

12. Miyazaki, Y., Oshima, K., Fogo, A., Hogan, B. L. M. & Ichikawa, I. Bone morphogenetic protein 4 regulates the budding site and elongation of the mouse ureter. Journal of Clinical Investigation 105, 863–873 (2000).

13. Combes, A. N. et al. Single cell analysis of the developing mouse kidney provides deeper insight into marker gene expression and ligand-receptor crosstalk (Development, (2019) 146, 12, 10.1242/dev.178673). Development (Cambridge) 146, (2019).

14. Lennartz, M. et al. Cadherin-16 (CDH16) immunohistochemistry: a useful diagnostic tool for renal cell carcinoma and papillary carcinomas of the thyroid. Sci Rep 13, 1–12 (2023).

15. Wertz, K. & Herrmann, B. G. Kidney-specific cadherin (cdh16) is expressed in embryonic kidney, lung, and sex ducts. Mech Dev 84, 185–188 (1999).

16. Prozialeck, W. C., Lamar, P. C. & Appelt, D. M. Differential expression of E-cadherin, N-cadherin and beta-catenin in proximal and distal segments of the rat nephron. BMC Physiol 4, 1–36 (2004).

17. Little, M., Georgas, K., Pennisi, D. & Wilkinson, L. Kidney development: Two tales of tubulogenesis. Curr Top Dev Biol 90, 193–229 (2010).

18. Wu, H. et al. Comparative Analysis and Refinement of Human PSC-Derived Kidney Organoid Differentiation with Single-Cell Transcriptomics. Cell Stem Cell 23, 869–881.e8 (2018).

19. Jung, J. Y. et al. Expression of epidermal growth factor in the developing rat kidney. Am J Physiol Renal Physiol 288, 227–235 (2005).

20. Kaiser, E., Kuzmits, R., Pregant, P. & Worofka, W. Clinical biochemistry of neuron specific enolase. Clinica Chimica Acta 183, 13–32 (1989).

21. Magella, B. et al. Cross-platform single cell analysis of kidney development shows stromal cells express Gdnf. Dev Biol 434, 36–47 (2018).

22. Rogakou, E. P., Pilch, D. R., Orr, A. H., Ivanova, V. S. & Bonner, W. M. DNA double-stranded breaks induce histone H2AX phosphorylation on serine 139. Journal of Biological Chemistry 273, 5858–5868 (1998).

23. Uchimura, K., Wu, H., Yoshimura, Y. & Humphreys, B. D. Human Pluripotent Stem Cell-Derived Kidney Organoids with Improved Collecting Duct Maturation and Injury Modeling. Cell Rep 33, 108514 (2020).

24. Montesano, R., Matsumoto, K., Nakamura, T. & Orci, L. Identification of a fibroblast-derived epithelial morphogen as hepatocyte growth factor. Cell 67, 901–908 (1991).

25. Desgrange, A. et al. HNF1B controls epithelial organization and cell polarity during ureteric bud branching and collecting duct morphogenesis. Development (Cambridge) 144, 4704–4719 (2017).

26. Kumar, S. V. et al. Kidney micro-organoids in suspension culture as a scalable source of human pluripotent stem cell-derived kidney cells. Development (Cambridge) 146, (2019).

27. Gerdes, J. et al. Cell cycle analysis of a cell proliferation-associated human nuclear antigen defined by the monoclonal antibody Ki-67. The Journal of Immunology 133, 1710–1715 (1984).

28. Takahashi, K. & Yamanaka, S. Induction of Pluripotent Stem Cells from Mouse Embryonic and Adult Fibroblast Cultures by Defined Factors. Cell 126, 663–676 (2006).

29. Li, J. et al. Roles of Krüppel-like factor 5 in kidney disease. J Cell Mol Med 25, 2342–2355 (2021).

30. Bohnenpoll, T. & Kispert, A. Ureter growth and differentiation. Semin Cell Dev Biol 36, 21–30 (2014).

31. Schwab, K. et al. A catalogue of gene expression in the developing kidney. Kidney Int 64, 1588–1604 (2003).

32. Tan, R. J. & Liu, Y. Matrix metalloproteinases in kidney homeostasis and diseases. Am J Physiol Renal Physiol 302, (2012).

33. Terzic, J., Muller, C., Gajovic, S. & Saraga-babic, M. Expression of PAX2 gene during human development. 707, 701–707 (1998).

34. Kaku, Y., Taguchi, A., Tanigawa, S., Haque, F. & Sakuma, T. PAX2 is dispensable for in vitro nephron formation from human induced pluripotent stem cells. 1–12 (2017) doi:10.1038/s41598-017-04813-3.

35. Buhl, E. M. et al. Dysregulated mesenchymal PDGFR-b drives kidney fibrosis. 1–20 (2020) doi:10.15252/emmm.201911021.

36. Dalghi, M. G. et al. Expression and distribution of PIEZO1 in the mouse urinary tract. Am J Physiol Renal Physiol 317, F303–F321 (2019).

37. Svenningsen, P., Hinrichs, G. R. & Zachar, R. Physiology and pathophysiology of the plasminogen system in the kidney. 1415–1423 (2017) doi:10.1007/s00424-017-2014-y.

38. Lin, Y., Wang, H., Huang, D., Hsieh, P. & Lin, M. Neuronal Splicing Regulator RBFOX3 (NeuN) Regulates Adult Hippocampal Neurogenesis and Synaptogenesis. 3, 1–17 (2016).

39. Lee, J. M., Dedhar, S., Kalluri, R. & Thompson, E. W. The epithelial – mesenchymal transition: new insights in signaling, development, and disease. 172, 973–981 (2006).

40. Sazonova, O. et al. Stanniocalcin-1 secretion and receptor regulation in kidney cells. Am J Physiol Renal Physiol 294, 788–794 (2008).

41. Czerniecki, S. M. et al. High-Throughput Screening Enhances Kidney Organoid Differentiation from Human Pluripotent Stem Cells and Enables Automated Multidimensional Phenotyping Resource. Stem Cell 22, 929–940.e4 (2018).

42. England, A. R. et al. Identification and characterization of cellular heterogeneity within the developing renal interstitium. Development 147, (2020).

43. Selmabel, E., Biology, C. & Haven, N. The Tight Junction Protein ZO-1 Is Concentrated Along Slit Diaphragms of the Glomerular Epithelium. J Cell Biol 11, 1255–1263 (1990).

